# Rational Design of Multiclade Coronavirus Spike Immunodominant Domain Nanoparticles to Elicit Broad Antibody Responses

**DOI:** 10.1101/2025.10.01.679649

**Authors:** Christian K.O. Dzuvor, Sydney Moak, Lindsay R. McManus, Abigail Thomas, Abigail E. Dzordzorme, Taewoo Kim, Jeswin Joseph, Valerie Foley, Ingelise J. Gordon, Laura Novik, LaSonji A. Holman, Lesia K. Dropulic, Ryan P. McNamara, Kizzmekia S. Corbett-Helaire

## Abstract

Four seasonal endemic human coronaviruses (EhCoVs), HKU1-CoV, OC43-CoV, 229E-CoV, and NL63-CoV, are culprits of mild upper respiratory and periodic severe diseases in vulnerable populations. Despite their prevalence, understanding EhCoVs’ antigenic and immune signatures remains elusive. SARS-CoV-2 has evolved as the fifth EhCoV, requiring seasonal vaccination in most parts of the world, and currently, no other EhCoV vaccines are available. SARS-CoV-2 co-infection with EhCoVs increases disease severity; thus, combined vaccination may provide increased protection against seasonal EhCoVs overall. Here, we explored Spike (S) receptor binding domain (RBD) vs. N-terminal domain (NTD) B-cell immunodominance in EhCoV-positive convalescent donors and immunogenicity in mice. We found that while antibody and B-cell isotypes were relatively dominant to S NTD, mice immunized with S RBD elicited significantly higher binding and neutralizing antibody (nAb) responses. With that knowledge, we used computational methods to infer that EhCoV S sequences evolve into two main clades and designed chimeric immunodominant domains (IDDs) from both clades for each EhCoV. IDDs were scaffolded onto two-component nanoparticles (NPs) displaying each IDD separately (monovalent IDD NP); three ß-EhCoV IDDs (Mosaic-3 IDD NP); or five EhCoVs IDDs (Mosaic-5 IDD NP). Mice immunized with mosaic IDD NPs, but not soluble IDD antigens nor monovalent IDD NPs, elicited potent, broadly cross-reactive binding and neutralizing antibody (Ab) responses against SARS-CoV-2 variants, other EhCoVs, and Sarbecoviruses. System serology revealed that all four IDD immunogens elicited distinct Ab subclasses and Fc-effector functions, with mosaic-5 IDD NPs eliciting the most *de novo* Ab subclasses, distributions, and broader Fc-mediated immune mechanisms. Dissection of vaccine-immune sera revealed polyclonal Ab responses against multiple non-overlapping cross-reactive S epitopes. Due to elicitation of broad Ab responses with combinatory functionality, IDD NPs open new horizons for developing first-in-class supraseasonal EhCoV vaccine candidates, with potential to decrease frequent SARS-CoV-2 sequence updates and protect against other EhCoVs. Moreover, elicitation of Ab breadth that spans pandemic-threat Sarbecoviruses gives mosaic IDD NPs promise towards pandemic preparedness.

## Main

Coronaviruses (CoVs) are zoonotic threats poised for human emergence, as most recently exemplified by severe acute respiratory syndrome coronavirus 2 (SARS-CoV-2), the causative agent of COVID-19^1^. To date, seven human CoVs are known to cause respiratory diseases of varying severity. Three (SARS-CoV, MERS-CoV, SARS-CoV-2) are highly pathogenic and deadly. The other four (HKU1-CoV, OC43-CoV, 229E-CoV, and NL63-CoV), classified as endemic human CoVs (EhCoVs), are culprits of up to 30% of “common colds” in any given “flu season”^2^. Although primarily associated with mild symptomatic illness and upper respiratory infection, sometimes EhCoV infection causes severe lower respiratory disease with occasional fatal outcomes, primarily in high-risk populations such as infants, the elderly and immunocompromised individuals^3,4^. Like SARS-CoV-2, EhCoV disease burden falls disproportionately on the elderly with specific comorbidities and immunocompromised patients^3^.

Despite frequent illness and/or reinfections every 6-8 months, no licensed EhCoV prophylactics, therapeutics, or vaccines exist. EhCoVs portend the future of SARS-CoV-2, and as SARS-CoV-2 has transitioned into the fifth seasonal EhCoV, disease burden from all EhCoVs has increased^5–7^. Additionally, EhCoV co-infection with SARS-CoV-2 presents a greater risk for severe respiratory illness and increased hospitalization rate^8–10^. Numerous licensed COVID-19 vaccines are available and highly effective in reducing hospitalization and severe disease^11–13^. Yet, no licensed COVID-19 vaccine with pan-EhCoV protective immunity is available. COVID-19 vaccine-induced immunity is predominantly provided by neutralizing antibodies (nAbs) targeting spike (S) protein, a homotrimeric complex of two functional subunits, S1 and S2, that facilitate receptor binding and membrane fusion, respectively. Nevertheless, frequent vaccine updates are required due to the emergence of SARS-CoV-2 variants and waning vaccine-induced immunity^14–16^. Also, no licensed COVID-19 vaccine with pan-EhCoV protective immunity is currently available. Indeed, several studies have demonstrated that COVID-19 vaccine-induced immunity is not associated with cross-reactive neutralization and protection against the other four EhCoVs^17–19^. Conversely, EhCoV antibodies (Abs) in previously infected humans do not protect against SARS-CoV-2 ^20,21^, neither do immune responses induced by EhCoV S vaccination^22^. Therefore, combining vaccination against other EhCoVs with the inevitable seasonal COVID-19 vaccination, may provide increased seasonal CoV protection^5,23^. Previous studies have demonstrated that SARS-CoV-2 infection boosts cross-reactive Abs to EhCoVs^17^. While boosted Abs primarily target the better-conserved S2 subunit, they were not associated with neutralizing activity^17^ or protection^24^.

To assess the feasibility of developing pan-EhCoV vaccines, it is imperative to better understand the antigenic and/or immunogenic signatures among different EhCoV S proteins and their domains. Such understanding is essential to define which S domains should be included in pan-EhCoV vaccines to optimize broad protective immunity.

The S1 subunit contains an N-terminal domain (NTD), a C-terminal domain (CTD)/receptor-binding domain (RBD), and two subdomains, SD1 and SD2. EhCoVs can be further classified by their genera: β-hCoVs (SARS-CoV-2, HKU1-CoV and OC43-CoV) and α-hCoVs (229E-CoV and NL63-CoV). Pre-fusion S structures unveil stark architectural and conformational differences between pandemic-threat β-hCoVs (SARS-CoV, MERS-CoV, and SARS-CoV-2), endemic β-hCoVs (HKU1-CoV and OC43-CoV), and endemic α-hCoVs (229E-CoV and NL63-CoV), such as distinct packing configuration between NTDs and CTDs/RBDs. For instance, endemic β-CoV S1-CTDs are static as opposed to dynamically flipping like pandemic-threat β-hCoVs^25–27^. Due to these structural and conformational differences, we hypothesize that α- and β-EhCoVs S proteins will exhibit differing antigenicity and immunodominance (ID). ID is a phenomenon of hierarchical immunogenicity across domains or epitopes on the same antigen.

Here, using HKU1-CoV as a prototype, we explore Ab and B-cell immunodominance in sera and peripheral blood mononuclear cells (PBMCs) from convalescent donors who acquired infection with EhCoVs (EhCoV-positive) prior to and during the COVID-19 pandemic and evaluate domain immunogenicity at S1 subdomain resolution. We found that Ab and B-cell isotypes were relatively dominant to NTD in humans. Vice versa, mice immunized with RBD but not NTD elicited significantly higher binding and robust neutralizing responses. Therefore, RBD is termed herein as the immunodominant domain (IDD) for the β-hCoVs (SARS-CoV-2, HKU1-CoV and OC43-CoV). NTD was selected as IDD for α-hCoVs(229E-CoV and NL63-CoV) based on previously reported antigenic mapping^28^. Prior to vaccine design, we performed a phylogenetic reconstruction of S protein sequences from the five EhCoVs, including SARS-CoV-2. We observed similar antigenic drift among all five EhCoVs. Each EhCoV bifurcates immediately after the root and splits into two main clades, A and B. Based on this analysis, we generated chimeric IDDs for pan-EhCoV vaccine designs by permutation of IDDs from both clades. Next, chimeric IDDs were fused to a trimerization motif to enhance cross-reactive immunogenicity within each EhCoV subtype. For SARS-CoV-2, we patched the IDD trimer with additional lineage mutations to expand its cross-variant antigenic coverage. The resulting IDD trimers were scaffolded on two-component I53-50 NPs^29,30^. We utilized this vaccine design strategy to generate NP immunogens displaying one IDD trimer (IDD NP), three β-EhCoVs IDD trimers (Mosaic-3 IDD NP), and five EhCoVs IDD trimers (Mosaic-5 IDD NP). Compared to IDD trimers and NPs, mosaic IDD NPs were more immunogenic in mice, eliciting superior cross-reactive binding and neutralizing Ab responses and Fc-mediated effector functions. Our work describes pan-EhCoV vaccine designs with potential to replace conventional SARS-CoV-2 vaccines – a salient goal to reduce overall seasonal EhCoV disease burden.

### B-cell and Antibody immunodominance (ID) hierarchy following EhCoV Infection

Understanding immunogenic signatures of the four EhCoV and SARS-CoV-2 S proteins is imperative for target identification and effective pan-EhCoV vaccine development (**Fig. 1A**). We used HKU1-CoV as a representative to investigate hierarchical focusing of immune responses at S1 subdomain resolution. We generated HKU1-CoV NTD and RBD probes to quantify serum Ab responses and B-cells expressing antibody isotypes (IgG, IgD and IgM) in peripheral blood mononuclear cells (PBMCs) from five EhCoV-positive convalescent donors (**Fig. 1B, Supplementary Table 1**). First, we assessed antigenicity of the probes using our recently isolated 14 monoclonal Abs (mAbs) from one of the EhCoV-CoV-positive convalescent donors (Wang, *et al*., submitted). The 11 RBD-specific mAbs bound to the HKU1-CoV RBD probe (**Supplementary Fig. 1A**); whereas the two NTD-specific mAbs interacted with the NTD probe (**Supplementary Fig. 1B**), and neither probe bound to the S2-specific mAb (**Supplementary Fig. 1C**), together confirming the antigenic integrity of the probes (**Supplementary Fig. 1)**. Then, we assessed whether polyclonal Abs from the convalescent sera would react to the HKU1-CoV NTD and RBD probes by ELISA. Serum Abs from all donors reacted to HKU1-CoV NTD and RBD probes (**Fig. 1C**).

**Fig. 1:**
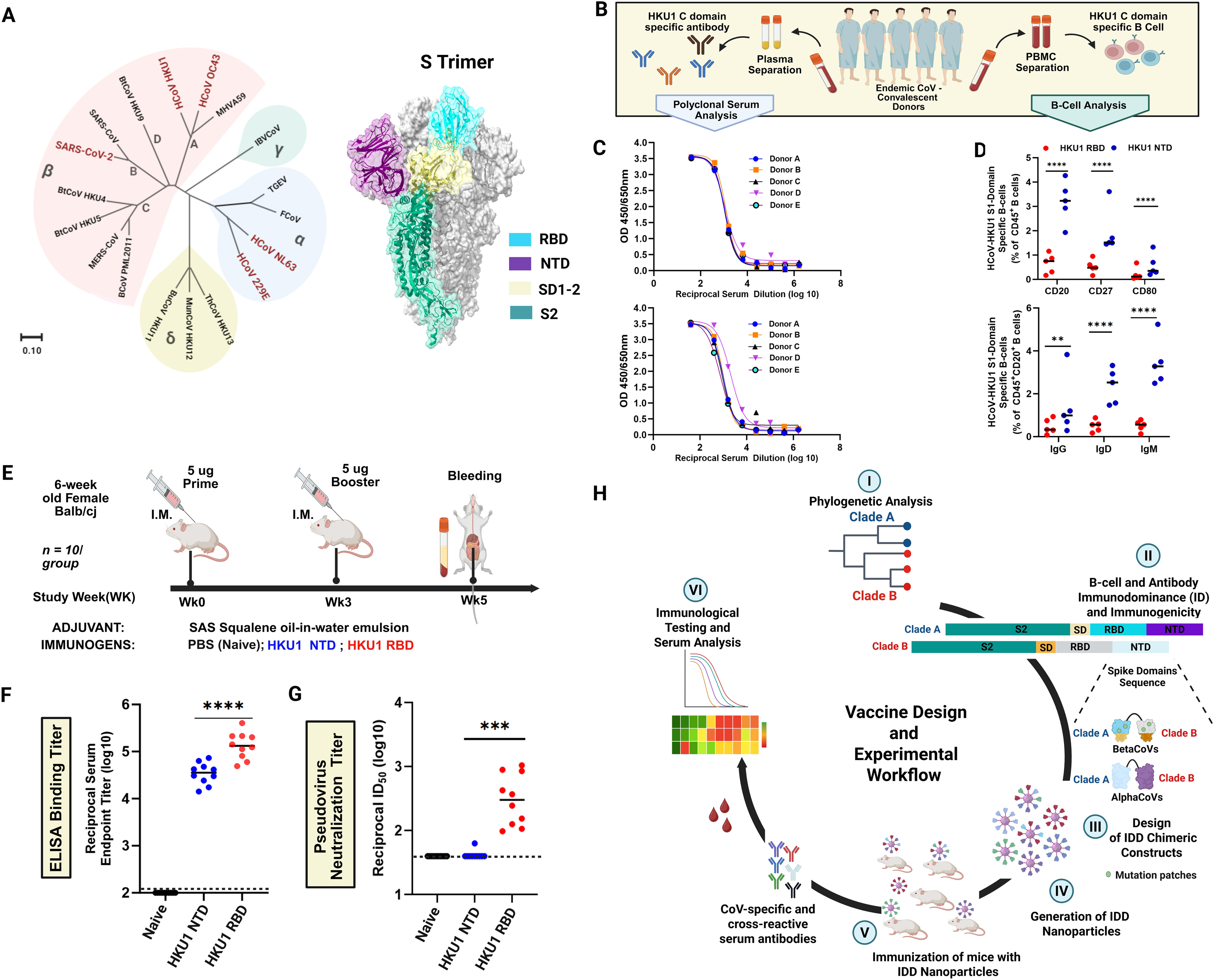
HKU1-CoV spike S1 domain-specific B-cell immunodominance (ID) and immunogenicity enable multiclade endemic human coronavirus (EhCoV) vaccine design. **(A)** Phylogenetic tree (left) of selected spike (S) protein sequences from CoV genera [alpha(α) beta(β), delta(δ) and gamma(γ)]; the five EhCoVs are highlighted in red. Structure model of S (right); RBD, NTD, SD1-2, and S2 are coloured in cyan, purple, yellow, and green, respectively. **(B)** Experimental workflow for HKU1-CoV-specific B-cell and antibody immunodominance. Convalescent donor sera were collected for serological analysis. Peripheral blood mononuclear cells (PBMCs) were used for B-cell analysis. **(C)** Sera were screened for binding by ELISA to HKU1-CoV NTD (upper) and RBD (lower) probes. **(D)** Identification and frequency of HKU1-CoV RBD- (red) and NTD-specific (blue) probe-specific B-cell subsets (top) and Ig isotypes (bottom). Number indicates percentage of gated cells. (E) Schematic illustration of mouse immunization schedule. Six-week-old female BALB/cJ (N=10/group) were immunized at weeks 0 and 3 with 5 µg of HKU1-CoV RBD (red) and NTD (blue) with SAS adjuvant and bled at week 5 for serology. Control mice(Naïve) were immunized with adjuvanted PBS (black). **(F)** Sera were screened for binding to HKU1-CoV S-2P by enzyme-linked immunosorbent assay (ELISA). **(G)** Sera were then assessed for neutralizing activity against HKU1-CoV pseudotyped virus, as shown by inhibitory dilution (ID50) to achieve 50% neutralization. The horizontal dashed line represents the limit of detection (LOD). **(H)** Vaccine design, concept, and experimental workflow. First, phylogenetic analysis of S sequences of each EhCoVs was performed **(I)**. Next, immunodominant domains (IDD) from selected spikes were fused, and NP constructs are designed (II-III). NPs were generated (IV). Mice are immunized with respective vaccines and bled (V). Sera were evaluated for cross-reactive antibody responses (VI). **(D, F-G)** Two–way ANOVA with Tukey’s post-hoc test was performed. Statistical significance is displayed as follows: **p<0.01, ∗∗∗p < 0.001; ∗∗∗∗p < 0.0001.

Next, we assessed the magnitude of B-cell responses by staining PBMCs with a mixture of fluorescently labelled NTD and RBD probes (**Fig. 1D, Extended Data Fig. 1**). Flow cytometric characterization revealed a significantly higher frequency of HKU1-CoV NTD-specific CD45^+^ B-cell isotypes (CD20, CD27, and CD80) compared to RBD-specific isotypes (**Fig. 1D, upper panel**). Also, the percentage of class-switched [immunoglobulin G-positive(IgG^+^), IgD^+^ and IgM^+^] peripheral CD20^+^ NTD-specific B-cells were substantially higher than the percentage of RBD-specific B-cells observed (**Fig. 1D, lower panel**). These data suggest NTD-specific B-cells dominated humoral responses in five HKU1-positive convalescent donors, possibly due to HKU1-CoV RBD occlusion and inaccessibility in its “down” conformation; this is consistent with recent molecular and structural findings suggesting that RBD accessibility (or “up” conformation) is triggered by NTD binding to sialic acid^26,31–33^. Oppositely, S1 RBD has been demonstrated to be the bona fide immunodominant domain (IDD) for other β-CoVs (SARS-CoV, MERS-CoV, SARS-CoV-2)^34,35^ and accounts for most potent neutralizing activity upon infection^34,36^ and vaccination^37,38^.

On this basis, we compared immunogenicity of the HKU1-CoV NTD and RBD domains following vaccination in BALB/cJ mice. Groups of 10 mice were immunized intramuscularly (IM) at weeks 0 and 3 with Sigma Adjuvant System (SAS)-adjuvanted formulations containing 5 µg of NTD or RBD antigens (**Fig. 1E**). At two weeks post-boost, RBD-immunized mice elicited significantly higher HKU1-CoV S protien-specific binding Ab responses than NTD-immunized mice (**Fig. 1F**). We then evaluated sera neutralization potential using HKU1-CoV pseudoviruses. Similarly, we observed that sera from RBD-immunized mice elicited robust nAbs, with geometric mean titer - reciprocal half-maximal inhibition dilution (GMT ID_50_) ranging from 2.1 to 3.2 logs. In contrast, NTD-immune sera did not induce detectable levels of S-specific nAbs (GMT ID_50_ = 1.6) (**Fig. 1G**). Similarly, in an accompanying submitted manuscript, we isolated a panel of HKU1-specific monoclonal antibodies (mAbs) using B-cells from one of the five HKU1-CoV-positive convalescent donors; we found RBD/CTD-specific mAbs have robust HKU1-CoV neutralizing activity. Conversely, NTD and S2-specific mAbs failed to neutralize HKU1-CoV pseudoviruses, in TMPRSS2-overexpressing cells^39^ (Wang, *et al*., submitted).

These immunological findings, together with structural insights^17,18,24,31,35,40^, highlight the importance of targeting neutralization-sensitive domains in next-generation pan-CoVs vaccine development^41,42^, particularly as we move towards seasonal COVID-19 vaccination in most parts of the world. To that end, we designed a rational vaccine concept, focusing on immunodominant domains (IDDs) for the five seasonal endemic-hCoVs^5^ (**Fig. 1H**).

### Immunodominant Domain (IDD) immunogen design, nanoparticle display, and characterization

Similar to influenza and other endemic human viruses^43^, the five EhCoVs are under strong positive selection, indicative of antigenic drift, genetic heterogeneity, adaptive evolution, and selective immune pressure^43–46^. Analogously, we performed ancestral or evolutionary phylogenetic reconstructions using EhCoV S protein sequences spanning clinical isolates and field samples. The five EhCoVs’ phylogenetic trees shared similar ladder-like topologies with lengthy trunks and short terminal branches (**Supplementary Fig. 2A-E**), evolving into two main segments, referred herein as clades A and B. Next, we selected candidate S proteins with better cross-reactive potential from both clades for each EhCoV. For each clade within an EhCoV phylogenetic tree, we hypothesized that uniquely or centrally positioned S protein possesses higher sequence or solvent-exposed residue identities and may have superior antigenic surfaces to elicit broadly cross-reactive antibody responses.

Therefore, RBD and subdomains (SD1-SD2) from selected S proteins from β-EhCoVs (HKU1-CoV, OC43-CoV, and SARS-CoV-2) clades A and B were fused via a linker to make chimeric immunodominant domains (IDDs) (**Fig. 2A**). RBD was chosen for HKU1-CoV and OC43-CoV based on aforementioned findings (**Fig. 1F-G**). For SARS-CoV-2, we chose RBD as it is the bona fide IDD^35,36^. Due to waning immune responses and emerging variants, we patched selected SARS-CoV-2 RBDs with key additional lineage mutations to expand its antigenic coverage (**Supplementary Fig. 2F)**. We included subdomains, SD1-2, as they are averse to change, functionally relevant, contain dominant epitopes, proximal to RBD, and cold spots for broadly cross-reactive antibodies^47,48^. Indeed, broadly cross-reactive neutralizing SD1-specific antibodies have been identified and synergize with RBD-targeting Abs to protect against SARS-CoV-2^47^. For α-hCoVs (229E-CoV and NL63-CoV), NTD from selected S proteins for both clades were fused to make a chimeric IDD (**Fig. 2A**), as a recent study found 229E-CoV NTD to be antigenically dominant and contain an antigenic supersite conserved in the NL63-CoV S protein^28^. Also, compared to 229E-CoV RBD, S protein trimer and intact monomeric S1 subunit induce more efficient nAbs against 229E-CoV^49^. Besides, NTD-targeting Abs have been shown to confer protection via other mechanisms such as Fc-mediated effector functions^50,51^. The chimeric IDD antigens reveal markedly distinct structures (**Supplementary Fig. 3A-C**).

**Fig. 2:**
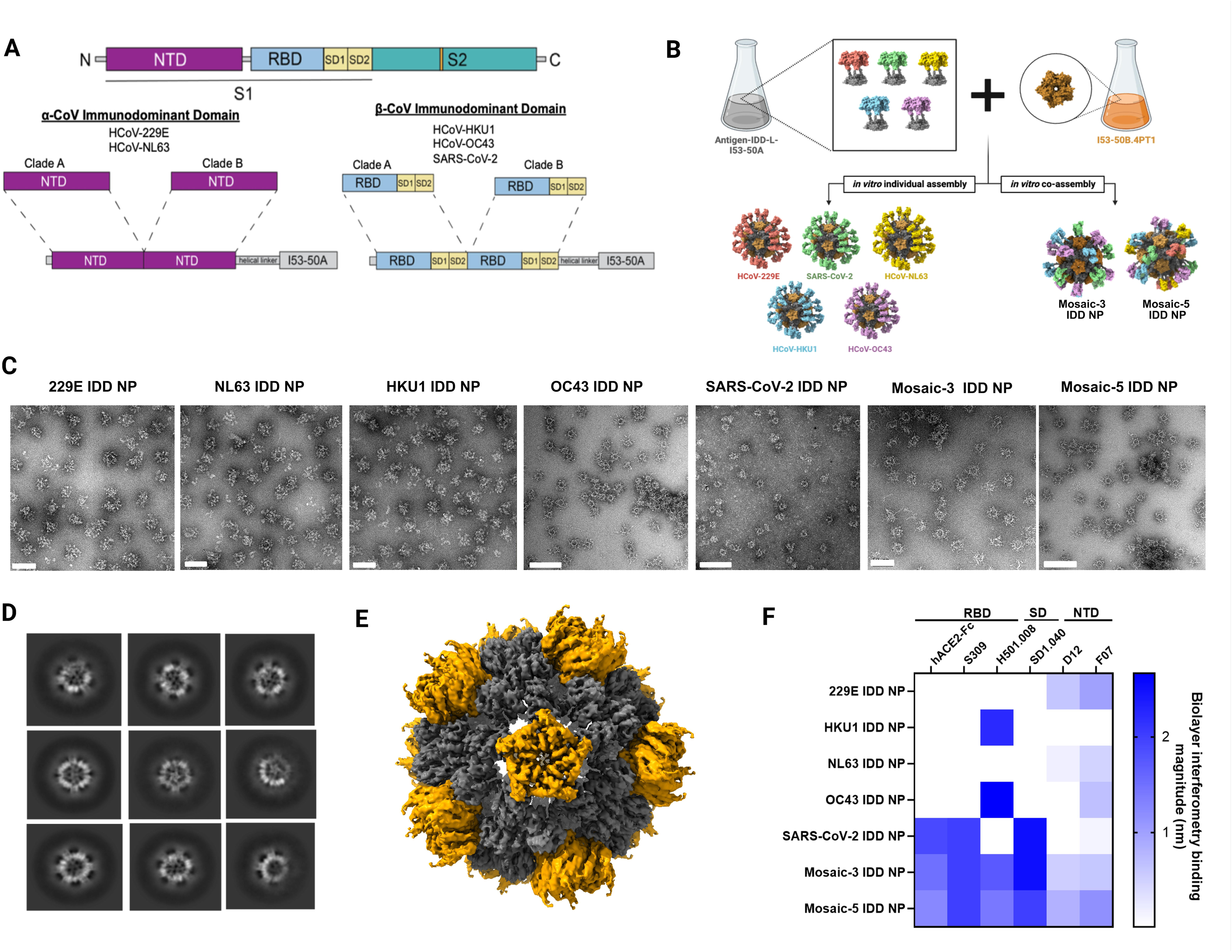
Design and characterization of IDD and IDD NP immunogens. **(A)** Schematic diagram of CoV S protein domain architecture (upper) and chimeric immunodominant domain design (lower) for endemic α-hCoVs (229E and NL63) and β-hCoVs (HKU1, OC43, and SARS-CoV-2). Two NTD or RBD-SD1-2 domains from clade A and B are dimerized. The C-terminal of the chimeric construct is fused to the N-termini of the NP trimer (I53-50A.1NT1) via a linker. **(B)** Schematic of in vitro assembly and NP formulation process. **(C)** Negative stain electron micrographs (EM) of purified NP immunogens. Scale bar, 50 nm. **(D)** 2D class averages of mosaic-5 IDD NP. **(E)** Single-particle cryo-electron microscopy reconstruction of mosaic-5 IDD NP, determined at 3.92 Å resolution. Cryo-EM density map was fitted with an atomic model (PDB:6P6F) and colored according to subunits in B. **(F)** Antigenic characterization of NP immunogens by bio-layer interferometry. Antigenicity was performed by assessing binding of immunogens to receptor, hACE2-Fc and S1 domain-specific mAbs. hACE2-Fc and S309 bind to SARS-CoV-2 RBD. H501.008 binds to HKU1-CoV and OC43-CoV RBD. SD1.040 binds to SARS-CoV-2 ubdomain, SD1. D12 and F07 bind to 229E-CoV and NL63-CoV NTD. Blue intensity signifies magnitude of binding.

As a target of nAbs, RBD immunogenicity can be augmented by multivalent display on NPs^52–54^, and mosaic display of heterologous RBDs can provide an avidity advantage for elicitation of superior cross-reactive B-cells and broad Ab responses^52,55,56^. Therefore, we generated immunogens that controllably co-display EhCoV IDDs using a computationally designed two-component protein NP, I53-50^29,30,57^. This system allows us to fine-tune and control valency of IDD antigens arrayed on the NPs. We genetically fused chimeric IDD antigens from the five EhCoVs to the N-terminus of I53-50A, the trimeric component of I53-50 NP, using a linker (**Supplementary Table 3**). Then, we expressed and purified each IDD-I53-50A protein via affinity and analytical size exclusion chromatography (SEC) (**Fig. 2B, Extended Data Fig. 2A-B).** The linker was used to ensure independent folding of the two components and enable optimal presentation and extension of IDD antigens from the NPs’ surfaces. Purified IDD-I53-50A trimers were mixed in equimolar amounts prior to the addition of purified pentameric I53-50B to generate mosaic NP immunogens that co-displayed three β-EhCoV IDDs (Mosaic-3 IDD-NP) or five EhCoV IDDs (Mosaic-5 IDD-NP) (**Fig. 2B**). In parallel, we produced NPs that solely displayed each IDD from the five EhCoVs. SEC revealed all proteins eluted at predominant peaks corresponding to the assembled icosahedral NPs (**Extended Data Fig. 2B**). Compared to IDD and bare I53-50 NP, IDD-NPs were monodisperse and homogeneous with intended icosahedral architecture by SEC, dynamic light scattering (DLS), and negative-stain transmission electron microscopy (NS-TEM) (**Fig. 2C-D, Extended Data Fig. 2B-C, Supplementary Fig. 3D-E**). We detected NP size and percent polydispersity (PDI) by DLS. The average hydrodynamic diameter for IDD-NP, mosaic-3 IDD-NP, and mosaic-5 IDD-NP were 44, 55 and 56.5 nm, respectively, compared to 17 nm for IDD trimers (**Extended Data Fig. 2C).** The PDIs of all NPs were less than 20%, indicating homogeneous NP size distribution. These results are further supported by a single particle cryo-electron microscopy reconstruction of mosaic-5 IDD-NP at a 3.92Å resolution (**Fig. 2E**, **Supplementary Fig. 4**). While the displayed IDD antigens were apparent upon imaging negatively stained and vitrified cryo-frozen NPs (**Fig. 2C-D, Supplementary Fig. 3E**), it was inadequately resolved after subsequent particle image averaging upon data processing (**Supplementary Fig. 4**). Nevertheless, fitting the computationally designed model, I53-50 into the density map supported the design’s precision and showed that genetically fusing IDD to I53-50A did not alter the structure of the two-component icosahedral NP (**Fig. 2E**).

Next, the NPs’ antigenic integrity was evaluated by bio-layer interferometry (BLI) using human ACE2, the cellular receptor for SARS-CoV-2, and EhCoV domain-specific mAbs. S309 and SD1.040, H501.008, and D12 and F07 bind to SARS-CoV-2, β-EhCoVs (HKU1-CoV and OC43-CoV) and α-EhCoVs (229E-CoV and NL63-CoV) respectively. 229E, HKU1, NL63, and OC43, and SARS-CoV-2 IDD and their IDD NPs bound to their respective mAbs (**Fig. 2F, Extended Data Fig. 2G**). Surprisingly, OC43 IDD and OC43 IDD NPs bound to F07, suggesting possible conserved epitopes between designed OC43-CoV IDD and cross-reactive α-EhCoVs NTD-targeting mAb (F07). Overall, mosaic −3 and −5 IDD-NPs bound to human ACE2 and all mAbs, whereas no single IDD or IDD-NP replicated this breadth of reactivity (**Fig. 2F, Extended Data Fig. 2D**). Intuitively, by blue intensity, mosaic-3 IDD NP bound with higher apparent affinity to hACE2-Fc, S309, SD1.040 and H501.008 compared to mosaic-5 IDD NP (**Fig. 2F**). Vice versa, mosaic −5 IDD NPs bound higher to D12 and F07. These higher affinity of mosaic −3 and −5 IDD NPs to the specific mAbs is attributed to the heterogeneity and valency of specific IDD antigens in each NP. Next, we tested ability of the immunogens to bind polyclonal Abs elicited by multiple EhCoV infections, allowing us to determine if the immunogens displayed Ab epitopes that are relevant to human infection, in turn alluding to the immunogens’ potential to amplify pre-existing antibody responses in humans. We performed ELISAs against the coated immunogens using convalescent sera acquired prior to and during the COVID-19 pandemic, which contained cross-reactive polyclonal Abs. Compared to bovine serum albumin (BSA), SARS-CoV-2 IDD, and SARS-CoV-2 IDD NP, we observed that all sera effectively reacted to the mosaic-3 and −5 NPs, and replicated a similar breadth of cross-reactivity (**Extended Data Fig. 3**)

### Elicitation of broadly cross-reactive antibody responses

To evaluate immunogenicity of IDD trimers and NPs, we vaccinated BALB/cJ mice with intramuscular injections of 5 µg of soluble IDD trimers, monovalent IDD I53-50 NPs, mosaic-3 IDD NPs, or mosaic-5 IDD NPs formulated with SAS, at weeks 0 and 3. Adjuvanted phosphate-buffered saline (PBS) was used as a control (Naïve group). Two-weeks post-boost, sera were collected for antibody assessments. Using ELISA, we measured serum antibody (IgG) binding titers against clade-specific CoV S proteins, matching the IDDs present in the vaccines (vaccine-matched) and contemporary CoV S proteins that were not present in the vaccines (vaccine-mismatched). Mice vaccinated with soluble IDD and monovalent IDD NPs exhibited less cross-reactivity **(Fig. 3A-B, Supplementary Fig. 5)**. For example, SARS-CoV-2 IDD NP vaccinated mice exhibited <4 logs geometric mean serum IgG endpoint titers to contemporary non-sarbecovirus S proteins (**Fig. 3B).** In contrast, mice vaccinated with mosaic-3 and −5 IDD NPs elicited robust cross-reactive Abs, in the range of 4 to 6 logs reciprocal endpoint titer, which reacted to all S protein except MERS-CoV S. Mosaic-3 and −5 IDD NPs elicited comparable cross-reactive Ab responses (**Fig. 3C-D)**. Radar plots more intuitively demonstrate that mosaic IDD NPs elicited the broadest Ab responses against S proteins from contemporary CoVs, clade A, and clade B EhCoVs, compared with SARS-2 IDD and IDD NPs (**Fig. 3E**). Only SARS-CoV-2 IDD-immune serum exhibited 3 log geometric mean IgG endpoint to MERS-CoV S protein. Despite RBD-sequence divergence between SARS-CoV-2 and MERS-CoV^58^, our rational inclusion of SD domains in our design may have accounted for this cross-reactive Ab response. The lack of binding by sera Abs from the remaining IDD NPs to MERS-CoV S protein could be due to occlusion of these epitopes when SARS-CoV-2 IDD is displayed on NP.

**Fig. 3:**
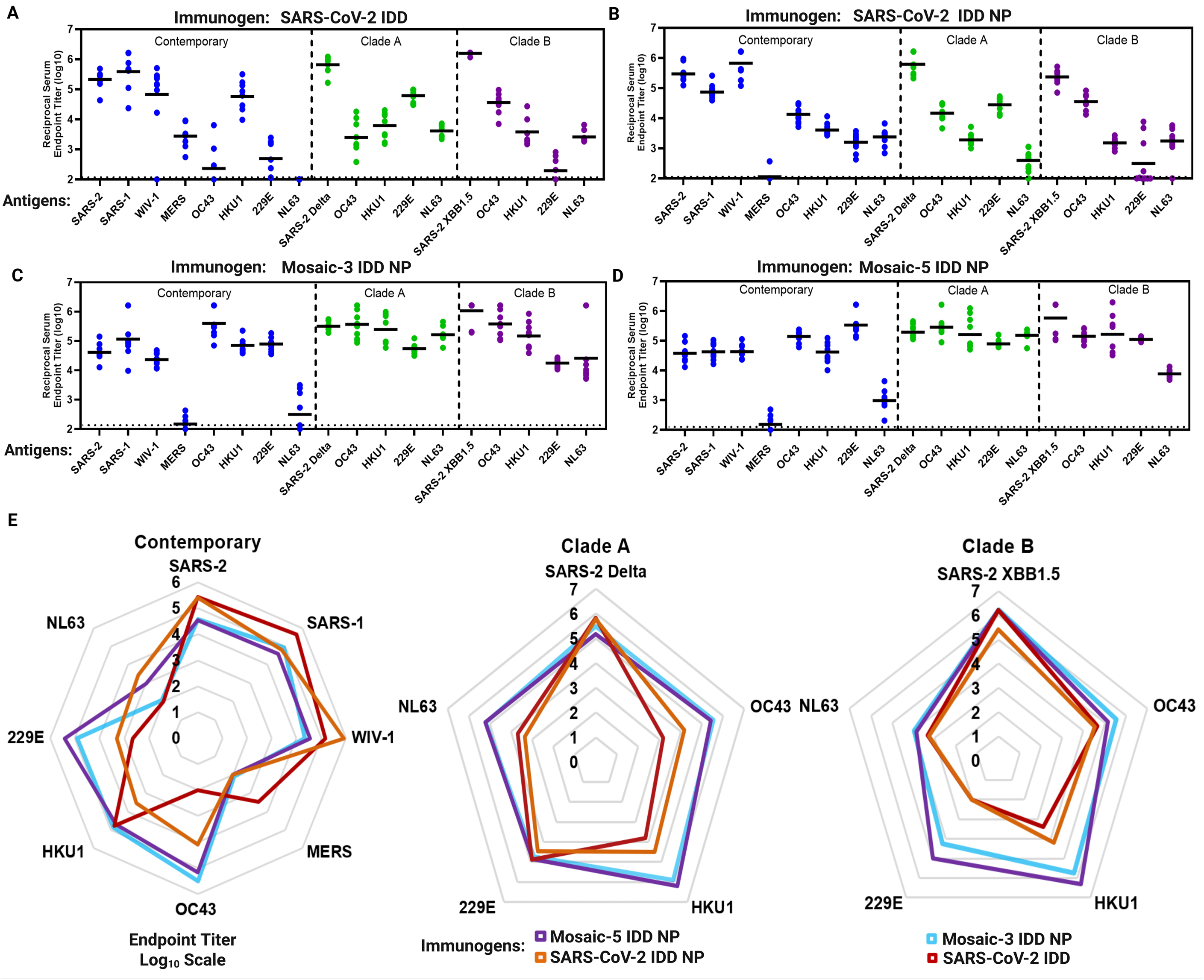
Cross-reactive antibody responses in SARS-CoV-2 IDD, SARS-CoV-2 IDD NP, Mosaic −3, and −5 IDD NP immunized mice. Female BALB/cJ mice (N = 10/group) were immunized at weeks 0 and 3 with 5 ug of SARS-2 IDD **(A)**, SARS-CoV-2 IDD NP **(B)**, Mosaic-3 IDD NP **(C)** and Mosaic-5 IDD NP **(D)** with SAS adjuvant and bled at week 5 for serology. Immune sera were assessed for antibody binding to various CoV S proteins by ELISA. Contemporary CoV S proteins are shown in blue. Clade A CoV S proteins are shown in green. Clade B CoV S proteins are shown in purple. Each dot represents an individual mouse. Horizontal dashed lines represent assay LOD. **(E)** Radar plot summarizing and comparing IgG serum titers of vaccinated mice against contemporary, clade A and Clade B CoV S proteins, from right to left. SARS-CoV 2 IDD = red, SARS-CoV-2 IDD NP = brown, Mosaic-3 IDD NP = blue, Mosaic-5 IDD NP – purple.

Taken together, these data demonstrate that our rationally designed IDDs are highly immunogenic against vaccine-matched and contemporary CoV strains compared to the naïve group (**Fig. 3, Supplementary Fig. 5**). Mosaic display of IDDs on NPs enhanced breadth of cross-reactive Ab responses to SARS-CoV-2 variants, SARS-CoV, WIV1-CoV, and contemporary, clade A, and B EhCoVs (**Fig. 3C-E**) – thus contributing to their promise as supraseasonal pan-EhCoV vaccines.

### Induction of broadly cross-neutralizing antibody responses

Given the ability of the SARS-CoV-2 IDD, SARS-CoV-2 IDD NP, mosaic-3 IDD NP, and mosaic-5 IDD NP to elicit robust SAR-CoV-2 WA1, Delta B. 1.617.2, and XBB1.5 S-specific binding Abs (**Fig. 3A-D**), we next assessed their ability to elicit nAbs against multiple SARS-CoV-2 variants of concerns (VOCs) (**Fig. 4A-B**). Serum from mice vaccinated with SARS-CoV-2 IDD NPs potently neutralized pseudoviruses bearing S proteins of all SARS-CoV-2 variants tested, with GMT ID_50_ ranging from 2.5 to 4 logs (**Fig. 4B, Extended Data Fig. 4**). Similarly, SARS-CoV-2 IDD immunization elicited equivalent nAb titers against all variants, except for Beta B.135.1; in which case, SARS-CoV-2 IDD NP elicited significantly higher B.135.1-specific nAbs (**Extended Data Fig. 4B**). Interestingly, serum from mice immunized with SARS-CoV-2 IDD and SARS-CoV-2 IDD NP neutralized contemporary and clade-specific NL63-CoV pseudoviruses, with a GMT ID50 titer of > 2.5 logs (**Fig. 4B, Extended Data Fig. 5E-F**). Additionally, SARS-CoV-2 IDD NP elicited nAbs against pandemic threat sarbecovirus, WIV1-CoV^59^ with GMT ID50 of 2.5 logs (**Fig. 4B, Extended Data Fig. 5G)**. Our SARS-CoV-2 vaccines are the first to demonstrate this breadth of CoV neutralization as this effect was not observed with currently approved COVID-19 vaccines^17,22^ or other COVID-19 vaccines in development^52,54,55,58^. Perhaps this notable result is attributed to SARS-CoV-2 and NL63 utilization of the same cellular receptor (ACE2) for entry. The result could also be due to rational inclusion of conserved subdomains 1-2 in our designs ^60^, further emphasizing the importance of rationally focusing immune responses onto broad immunogenic sites.

**Fig. 4:**
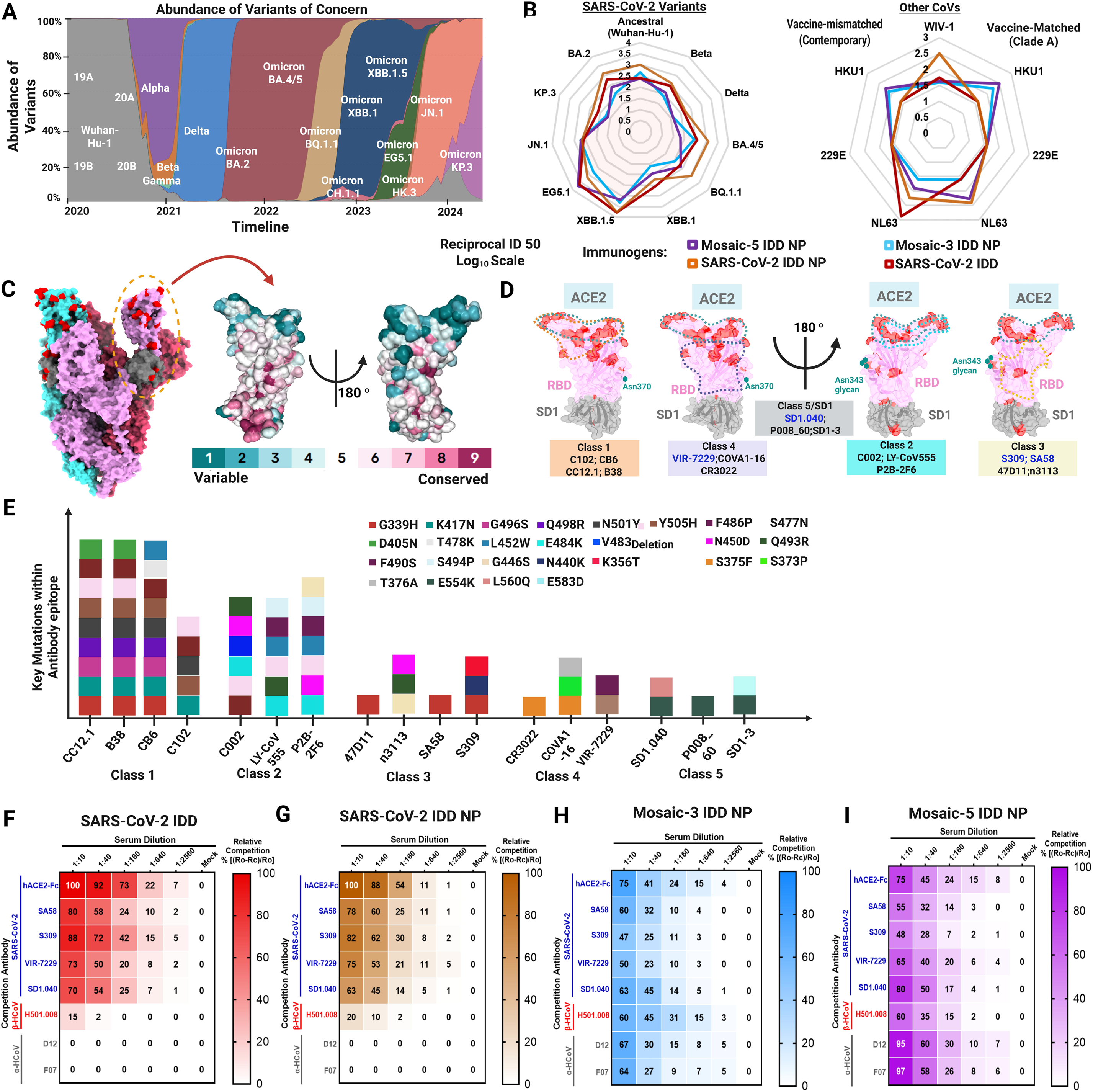
Cross-reactive neutralizing antibody responses in immunized mice against SARS-CoV-2 strains and EhCoVs. (**A**) Global prevalence and timeline of SARS-CoV-2 variant emergence. Data is from GISAID via CoVariants.org (**B**) Radar plots summarizing ID50 neutralization titers in SARS-CoV-2 IDD (red), SARS-CoV-2 IDD NP (brown), Mosaic −3 (blue), and −5 NP (purple) immunized mice against SARS-CoV-2 variants (left) and WIV-1, contemporary (mismatched), and matched EhCoVs (right). (**C**) Left, Structure of SARS-CoV-2 Spike glycoprotein (PDB: 6VXX), showing one “up” RBD with amino acid mutations found in variants highlighted in red. Right, Sequence conservation of SARS-CoV-2 variants calculated by ConSurf Database and overlaid on a surface representation of the SARS-CoV-2 RBD-SD1 structure (PDB:8D48) identifies conserved epitopes or regions. The color gradient on the bottom contains nine total steps or scores. (**D**) Epitope and footprint of hACE2 and five classes of antibodies on SARS-CoV-2 RBD-SD1. Prototype RBD and SD1 are shown as pink and gray surfaces (PDB:8D48). Residues that mutated in SARS-CoV-2 variants to date are indicated as red patches. Footprints of hACE2 and five classes of Abs are outlined in colored dotted lines. The N-linked RBD residue 343 glycan is indicated in a green sphere. (**E**) Key SARS-CoV-2 variants’ amino acid mutations found in binding sites of five Abs classes. (**F-I**) Heatmaps representing the relative competition of pooled mouse sera with mAbs, as assessed by BLI. Results from (Ro − Rc) × 100/Ro represent competition levels. Ro represents the highest binding signals of mAb without serum. Rc represents the highest binding signals of mAb tested with sera from vaccinated mice. Dilution level or inhibition level is represented as colored gradient bar. Strongest inhibition by certain mAbs is displayed as deepest color, indicating more abundance serum Ab binding to a similar epitope.

Additionally, despite 1/3^rd^ or 1/5^th^ the valency of each IDD in the mosaic-3 and −5 IDD NPs, respectively, these NPs elicited broad serum nAbs against SARS-CoV-2 variants and contemporary and clade-specific EhCoVs with GMT ID_50_ titers on par with SARS-CoV-2 monovalent IDD and IDD NPs (**Fig. 4B, Extended Data Figs. 4 and 5)**. As expected, mosaic-3 and −5 IDD NP elicited nAbs against contemporary HKU1-CoV pseudoviruses with GMT ID_50_ of 2.5 logs; mosaic-3 also neutralized clade A HKU1-CoV (GMT ID_50_ = 2.5 logs) and neither neutralized 229E-CoV **(Fig. 4B, Extended Data Fig. 5A-G)**. The lack of EhCoV neutralizing activity against α-EhCoVs, suggests that the robust Ab responses elicited by the mosaic IDD NPs have orthogonal functions that are non-neutralizing. Nevertheless, these data exemplify that our chimeric IDD approach enhanced immunogenic coverage against SARS-CoV-2 variants and EhCoVs compared to historical data with current COVID-19 vaccines^61^. Additionally, multivalent display of IDDs on NPs allows for reduced valency of each antigen with markedly equivalent ability to elicit nAbs against SARS-CoV-2 variants, enhanced immunogenicity against WIV1-CoV and HKU1-CoV, and modest clade A NL63-CoV nAbs. While the motivation for this approach was to elicit broad immunogenicity against SARS-CoV-2 variants and the other four seasonal EhCoVs^5,23^, these data suggest that these vaccine candidates (SARS-CoV-2 IDD NP and mosaic −3 and −5 IDD NPs) have the potential to not only serve as pan-EhCoVs vaccines but also protect against related Sarbecoviruses^59,62^, as relayed by immunogenicity against WIV1-CoV.

### Molecular basis and epitope specificity of IDD vaccine-elicited antibodies

The robust Ab responses elicited by our IDD vaccines begged the question of what cross-reactive Ab epitopes the vaccines targeted. Therefore, we mapped specificity of polyclonal serum Ab responses elicited by our vaccines against well-characterized cross-reactive Abs using a BLI antibody competition assay (**Extended Data Fig. 6A**)^63^. As SARS-CoV-2 S protein antigenic landscape has been well-characterized, we first mapped mutations from SARS-CoV-2 variants onto the SARS-CoV-2 Wuhan-1 S protein (PDB: 6VXX) and RBD-SD1 (PDB:8D48) (**Fig. 4C**), highlighting epitope conservation with previously characterized mAbs)^64^. Based on the spatial overlap of viral mutations, Ab epitopes, and S protein binding mode, we grouped these Abs into five classes (**Fig. 4D**). RBD-targeting Abs can be grouped into four classes based on previous classification^65–67^; class 5 consists of conserved SD1-2 epitope-targeting Abs. Recent studies revealed that certain class 3 and 4 mAbs can effectively neutralize a wide range of β-coronaviruses, such as RaTG13, Pangolin (Guangdong) CoV, SARS-CoV, SARS-CoV-2, WIV1-CoV, and SHC014^68–71^. Notably, these Abs maintain their neutralizing ability against new SARS-CoV-2 VOCs^68,71^ and are resilient to epitope diversification^68^. Also, some cross-reactive SD1 mAbs potently neutralize SARS-CoV-2 variants^47,72^. Evidently, the epitopes of these Abs involved few mutations (**Fig. 4E**). We selected Abs from class 3 (SA58^73^, S309^70^), (VIR-7229^68^), and 5 (SD1.040^47^). hACE2-Fc was used to determine the presence of ACE2-binding site Abs. For the other four EhCoVs, little was known about their S protein antigenic landscapes. As there are currently no cross-reactive HKU1/OC43-specific Abs, we used our recently discovered one, H501-008, which we isolated and characterized from an HKU1-CoV convalescent donor 501 herein (**Supplementary Table 1**) in an accompanying manuscript (*Wang, et al*. submitted). For α-hCoVs (229E-CoV and NL63-CoV), recently published cross-reactive NTD-targeting Abs (D12 and F07)^28^ were used.

Next, anti-Penta-HIS (HIS1K) biosensors coated with either SARS-CoV-2, OC43-CoV, or 229E-COV IDD trimers were first incubated with serially diluted IDD-immune sera and then with each of the mAbs (**Extended Data Fig. 6A**). Our results showed that as serum concentration increased during primary binding, secondary mAb binding declined exemplifying individual epitope specificity within the polyclonal serum Abs (**Extended Data Fig. 6B-C**). In contrast, sera with little-to-no similar Abs showed almost no competitive effects as shown with SARS-CoV-2 IDD NP-immune sera competed with H501-008 (**Extended Data Fig. 6C, right**). All SARS-CoV-2 mAbs (hACE2-Fc, SA58, S309, VIR-7229, SD1.040) effectively competed with sera from all vaccine groups (SARS-CoV-2 IDD, SARS-CoV-2 IDD NPs, and mosaic-3 and −5 IDD NPs) in a concentration-dependent manner (**Fig. 4F-I, Extended Data Fig. 6B-C)).**

Next, we measured individual sera relative competitiveness at each dilution. In comparison to other vaccine groups, we observed that serum from SARS-CoV-2 IDD NP immunized mice elicited the most potent inhibition with the five SARS-CoV-2-specific Abs, indicating SARS-CoV-2 IDD scaffolded to NPs enhances and exposes those epitopes more than SARS-CoV-2 alone or SARS-CoV-2-inclusive mosaic NPs **(Extended Data Fig. 6D**). In the SARS-CoV-2 IDD NP group, the highest competitive inhibition percentages were observed with hACE2-Fc (93%) and S309 (84%), followed by SA58 and SD1.040 (83%), and lastly VIR-7229 (67%), suggesting gradually decreasing epitope exposure from RBD to the transmembrane proximal region. We also observed that sera from mosaic-3 and −5 IDD NP vaccinated mice effectively inhibited binding of hACE2-Fc and mAbs against all five EhCoVs, recapitulating breadth of cross-reactive binding and neutralizing capacity. For mosaic-3 IDD NP, immune sera competed with SARS-CoV-2 and ß-EhCoV-specific mAbs up to 60%, whereas for α-EhCoV mAbs, competition was equal to or below 30% (**Extended Fig. 6D**). As expected, due to the inclusion of 229E- and NL63-CoV IDDs, mosaic-5 IDD NP elicited Abs that competed with α-EhCoV mAbs at 95% (at 1:10 serum dilution) (**Fig. 4I**). Overall, these data indicate that SARS-CoV-2 IDD NP elicits the most robust SARS-CoV-2 cross-reactive Ab responses. Additionally, immunization with all our IDD vaccines elicited Abs targeting multiple non-overlapping cross-reactive epitopes, including conserved neutralizing and immunodominant sites for major potent class 3 and 4 RBD-targeting Abs^36^. Elicitation of these types of Abs alludes to the promise of our vaccines’ effectiveness against emerging SARS-CoV-2 escape mutants and variants. For mosaic IDD NPs, while increasing valency decreased exposure to SARS-CoV-2-specific epitopes, added EhCoVs increased overall EhCoV epitope breadth—further pointing to validity of these vaccines as supraseasonal EhCoV vaccine candidates.

### Stimulation of fc-mediated antibody effector mechanisms

Given the moderate levels of cross-nAb responses elicited by mosaic IDD NPs and the proportion of SARS-CoV-2-specific Ab populations ( SA58^73^, S309^70^, and VIR-7229^68^) targeting class 3 and 4 epitopes in sera from all vaccinated mice (**Fig. 4F-I, Extended Data Fig. 6D**), we next investigated the ability of our vaccines to elicit Abs with Fc-mediated effector functionality. NAbs targeting class 3 and 4 epitopes in SARS-CoV-2 RBD have been found to promote potent Fc-effector functions^68,74,75^. Our mosaic-5 IDD NPs include α-EhCoV S protein NTD domains; NTD-targeting neutralizing and non-neutralizing Abs have been shown to elicit Fc-mediated effector functions^50,51^ and synergize to confer protection against CoV infection^76^. Prior to evaluating contributions of Fc effector functions, we profiled vaccine-induced Abs in individual sera using a system serology assay^77,78^ (**Supplementary Fig. 7**). We determined levels for serum Abs that bind to S proteins from contemporary/vaccine-mismatched CoV strains (SARS-CoV-2, HKU1-CoV, OC43-CoV, SARS-CoV, WIV1-CoV, 229E-CoV, or NL63-CoV) and IDD proteins from vaccine-matched CoV strains (SARS-CoV-2, HKU1-CoV, OC43-CoV, 229E-CoV, or NL63-CoV). We also profiled the sera’s Ab isotypes and subclass specificity (IgG1, IgG2a, IgG3, IgM, or IgA) to the S proteins, as well as the Abs’ ability to interact with specific Fcγ receptors (FcγR IIb, FcγR III, or FcγR IV). Naïve sera and binding to Ebolavirus glycoprotein (EBOV GP) were used as reference and negative binding controls, respectively (**Fig. 5A, Extended Data Fig. 7, Supplementary Fig. 6**). Compared to naïve sera (**Fig. 5A, left),** immune sera from vaccinated mice contained higher total IgG, IgG1, and IgG2a Abs against contemporary and clade-specific proteins (**Fig. 5A, right**). Overall binding profiles showed broader Ab reactivity in the Mosaic-3 and Mosaic-5 IDD NP vaccinated mice, particularly for IgG1 and polyfunctional IgG2a. Similarly, FcγR-binding antibodies, both inhibitory (FcγR IIb) and activating (FcγR III and FcγR IV), showed enhanced binding to the CoV S proteins for all vaccinated groups compared to naïve. However, the Mosaic-3 and Mosaic-5 IDD NP again showed generally greater breadth of binding and higher titers compared to the SARS-CoV-2 IDD and SARS-CoV-2 IDD NP groups.

**Fig. 5.**
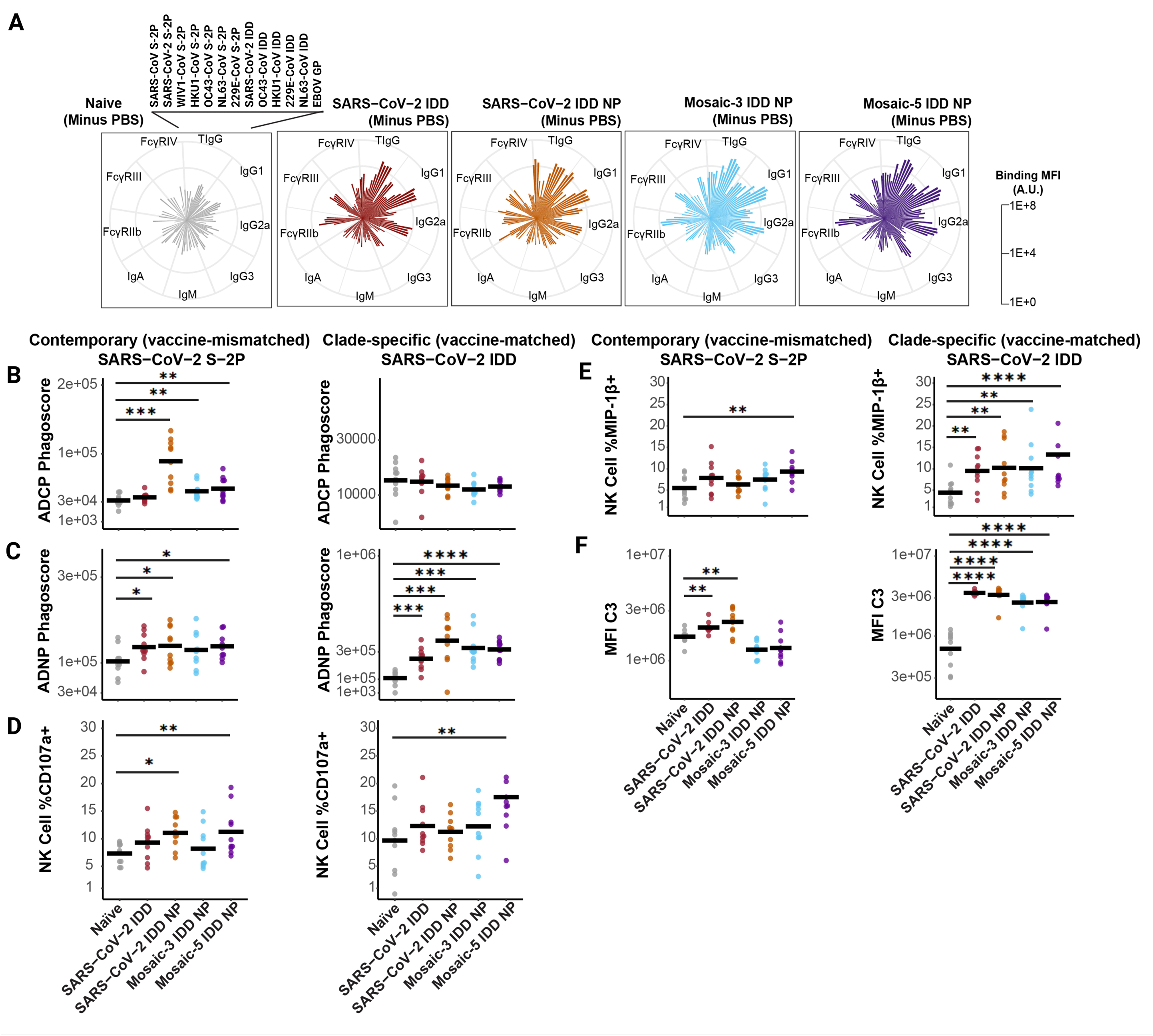
Systems serology analysis of IDD vaccine-induced immune sera. (**A**) Radar plots showing antibody binding and FcγR-binding levels to an array of CoV Spikes. Radars show isotype (IgG, IgA, IgM), subclass (IgG1, IgG2a, IgG3), and FcγR-binding antibodies for each treatment group minus background (PBS). Shown on the right side is the scaling. All values are in mean fluorescence intensity (MFI) in arbitrary units (AU). Ebolavirus glycoprotein (EBOV Gp) was used as an antigen specificity negative. (**B-F**) Fc-mediated effector functions of the naïve and immune sera after vaccination with IDD NPs against contemporary SARS-CoV-2 S-2P (left-hand columns) and clade-specific SARS-CoV-2 IDD protein (right-hand columns). Antibody-mediated cellular phagocytosis with monocytes (ADCP, **B**) or neutrophils (ADNP, **C**) using IDD-vaccine-induced immune and naïve sera and beads coated with SARS-CoV-2 S-2P and SARS-CoV-2 IDD proteins. **D-E**, Antibody-dependent natural killer cell activation (ADNKA) using IDD-vaccine-induced immune and naïve sera and beads coated with SARS-CoV-2 S-2P and SARS-CoV-2 IDD proteins. **D**, Percentages of NK cells expressing CD107a against the antigens by vaccine group. **E**, Percentages of NK cells expressing MIP-1β. The experiment was run with PBMCs from two separate donors. (**F**) Antibody-dependent complement deposition (ADCD) on beads coated with SARS-CoV-2 S-2P and SARS-CoV-2 IDD proteins and incubated with naïve or immune sera. In all figures, the horizontal black bar indicates the group mean value (N=10 mice per group). Each dot represents an individual mouse in a group. Two–way ANOVA with Tukey’s post-hoc test was performed. The statistical significance is displayed as follows: ∗p < 0.05; ∗∗p < 0.01; ∗∗∗p < 0.001; ∗∗∗∗p < 0.0001. Source data file is provided.

Given the distinctions observed in humoral profiles from the vaccinated groups, particularly among FcγR-binding antibodies, we next evaluated Fc-effector functions. We focused on antibody-dependent cellular phagocytosis by monocytes (ADCP), antibody-dependent neutrophil phagocytosis (ADNP), antibody-dependent NK cell activation (ADNKA), and antibody-dependent complement deposition (ADCD). Each mechanism was tested in response to the same contemporary and clade-specific proteins to evaluate the breadth of effector functions against different CoVs (**Fig. 5B, Extended Data Fig. 7, Supplementary Fig. 6**). Compared to naïve sera, vaccine-induced immune sera promoted greater ADNP and ADCP functionality against most contemporary and clade-specific antigens. ADCP to contemporary, vaccine-mismatched targets (SARS-CoV-2, SARS-CoV, WIV1-CoV) was present for SARS-CoV-2 IDD NP and mosaic-3 and mosaic-5 IDD NP vaccinated groups compared to naïve serum (**Fig. 5B**). ADNP trended similarly for both clade-specific vaccine-matched (SARS-CoV-2 S trimer) and contemporary targets for these groups (**Fig. 5C**). ADCP and ADNP responses to OC43 and HKU1 also showed enrichment for the vaccinated groups, with the mosaic-3 and mosaic-5 IDD NP vaccinated groups being most significantly pronounced, particularly for contemporary, vaccine-mismatched targets (**Figs. 5B-C, Extended Data Fig. 7A-B, Supplementary Fig. 6A-B**).

For ADNKA analysis, we measured percentages of NK cells expressing CD107a, a degranulation marker, and macrophage inflammatory protein 1 beta (MIP-1β), an NK cell activation marker and pro-inflammatory cytokine. For both vaccine-matched and vaccine-mismatched, CD107a responses were observed for the mosaic-5 IDD NP vaccinated group. The SARS-CoV-2 IDD NP vaccine group also showed CD107a responses to vaccine-mismatched SARS-CoV-2 S-2P (**Fig. 5D**). Likewise, for MIP-1β expression, the mosaic-5 IDD NP vaccine group showed NK responses against clade-specific and contemporary targets (**Fig. 5E**, left). For the vaccine-matched clade-specific response, all immunized groups showed significant MIP-1β stimulation compared to the naïve group (**Fig. 5E**, right). Neither immune sera enhanced ADNKA in response to HKU1-CoV or OC43-CoV S proteins (**Extended Data Fig. 7C-D**). Nevertheless, ADCD was significantly greater for immune sera from all vaccine groups in response to all clade-specific proteins compared to naïve sera. Only SARS-CoV-2 IDD and IDD NP immune sera significantly promoted ADCD against contemporary (SARS-CoV-2, HKU1-CoV, OC43-CoV, SARS-CoV and WIV1-CoV) S proteins.

Overall, these findings indicate that our vaccine sera contain diverse Abs against CoV proteins, capable of binding to FcγRs and inducing an array of Fc-mediated effector functions. Notably, mosaic-5 IDD NP generated some de novo Ab responses and showed a breadth of Fc-mediated immune protective mechanisms comparatively.

## Discussion

There are no approved vaccines, prophylactics, or therapeutics against the four “common cold” EhCoVs, which circulate seasonally typically causing mild respiratory tract infections and sometimes severe lower respiratory disease in vulnerable populations^3,17^. We are five seasons into repeated COVID-19 vaccinations due to waning immunity and SARS-CoV-2 VOC emergence, not considering increased disease burden caused by co-infection with other EhCoVs^8^. Thus, we are motivated by the prospect of a seasonal EhCoV vaccine that protects against all five EhCoVs and, importantly, withstands future SARS-CoV-2 variants—namely a “pan-EhCoV vaccine”. To that end, we set forth to design and characterize vaccine candidates towards two major goals: (1) elicitation of broad and robust Ab responses against contemporary and historical EhCoV strains and (2) elicitation of broad and robust Ab responses against vast SARS-CoV-2 VOCs.

S protein is the key antigenic target for CoV vaccine design^25^, and also most pan-sarbecovirus vaccine designs^42,79^ utilize RBD due to its ID and nAb landscape^36^. Our recent work emphasizes rational pan-CoV vaccine design must be guided by a comprehensive understanding of differences in S protein antigenic and immune landscapes of diverse CoV clades^38,80^. However, this understanding remains poorly defined, particularly for ß-EhCoVs, despite recent structural insights^17,18,24,31,35,40^. Specifically, before this study, little was known about ß-EhCoV B-cell and Ab ID. Herein, our primary aim was to define Ab and B-cell ID in EhCoV-positive convalescent donors and evaluate domain-specific immunogenicity. That allowed us to identify target domains, “immunodominant domains (IDDs)”, that we then rationally included in pan-EhCoV vaccine designs. While B-cell subsets and Ab isotypes were relatively dominant to NTD in EhCoV convalescent humans, mice immunized with HKU1-CoV RBD elicited significantly higher binding and neutralizing Ab responses compared to HKU1-CoV NTD-immunized mice. This knowledge enabled us to design multiclade IDD trimers and scaffold them to monovalent and mosaic NPs vaccines, rendering vaccines that elicited potent vaccine-matched and mismatched cross-reactive binding, neutralizing, and Fc-mediated antibody functionality against four EhCoVs (HKU1-CoV, OC43-CoV, 229E-CoV, and NL63-CoV), SARS-CoV-2 VOCs, and other distantly related Sarbecoviruses (SARS-CoV and WIV1-CoV).

These findings go beyond previous next-generation universal CoV vaccine concepts^41,42^, which elicit cross-reactive binding and neutralizing Ab responses with varied potency and breadth against SARS-CoV, SARS-CoV-2 and variants^54,58,81–83^ or against specific CoV subtypes^52,53,55,84^, but limited Fc-mediated effector functionality. Neutralizing Abs and Fc-mediated effector functions are independent immune correlates of protection against CoV infections in humans^51^. Vaccines, such as our SARS-CoV-2 IID NPs, that can induce both with a breadth of cross-reactivity would have advantages over approaches that induce one or the other, making them promising supraseasonal EhCoV vaccine candidates with potential to replace current and frequently updated seasonal COVID-19 vaccines. Moreover, the breadth and combinatory functionality of Abs elicited by our mosaic-3 and −5 NPs suggest they may be able to provide consistent immunity against seasonal EhCoVs and be effective in the event of outbreaks of antigenically mismatched Sarbecoviruses, such as RaTG13, SHC014, and pangolin coronavirus GXP4L^62^.

Further, our data revealed that our IDD vaccines elicited SARS-CoV-2 Ab responses against multiple non-overlapping RBD sites, targeted by potent cross-reactive SARS-CoV-2 Abs^36,71^. To date and to the best of our knowledge, no other vaccine concept has been shown to achieve this feat. Several SARS-CoV-2 isolates evade mAbs and immune serum recognition^85^, therefore mAb combinations have been evaluated and shown to overcome immune evasion^86,87^. In accordance, the diverse polyclonal Ab responses elicited by our IDD vaccines, targeting several distinct cross-reactive epitopes, could account for the breadth of cross-neutralizing activity observed. Thus, we anticipate immunity elicited by these vaccines should mitigate the likelihood of emergence of escape mutants^88^. As an additional benefit, such effects are likely to carry over to variants of other EhCoVs through vaccination with our mosaic IDD NPs, as demonstrated by breadth of Ab responses to contemporary and clade A/B EhCoV strains.

Finally, although investigating the structural basis for the breadth of cross-reactivity elicited by our vaccines is required, we showed that rationally focusing vaccine-elicited Ab responses onto complex immunogens requires deeper comprehension of Ab and B-cell ID. For EhCoVs, it remains elusive, but initial approaches to characterize B-cell and Ab ID hierarchies, like we did for convalescent EhCoV-positive humans, are vital. It will be critical to continue to understand how ID changes over time following each EhCoV infection (ie, in the acute vs. convalescent phase), how ID evolves over a lifespan following multiple EhCoV infections, and how ID varies with disease burden, comorbidities and age. Using our ID findings to inform rational vaccine design provides a framework for vaccine design built upon understanding of immunogenic signatures for not only EhCoVs but other highly variable viruses as well. Moreover, given that majority of people have previously been infected with multiple EhCoVs, and considering approximately 14 billion COVID-19 vaccine doses have been administered globally^89^, it will be important to assess our IDD vaccines’ ability to elicit broad antibody responses in the context of pre-existing immunity or original antigenic sin. Additionally, there is still a need to assess protective efficacy of these vaccines, but given current understanding of correlates of protection^51,90,91^, the broad, potent, and multiple Ab functions elicited by our IDD vaccines are predictive of protection against SARS-CoV-2 VOCs, EhCoVs, and, notably, pandemic-threat Sarbecoviruses.

## Methods

### EXPERIMENTAL MODEL AND SUBJECT DETAILS

#### Cell lines

HEK293T/17 (ATCC CRL-11268) cells and 293T cells stably overexpressing human ACE2 (provided by M. Farzan, Scripps Research Institute) were cultured in Dulbecco’s modified Eagle’s medium (DMEM) supplemented with 10% FBS, 2 mM glutamine, and 1% penicillin–streptomycin. MF_293T-hACE2^92^ was supplemented with 3 µg/mL puromycin. 293T-TMPRSS2 and 293T-ACE2-APN cells were cultured in DMEM supplemented with 4.5g/L glucose and sodium pyruvate without L-glutamine, 10% FBS, 1% penicillin-streptomycin, 1x Glutamine, 1x MEM Nonessential Amino Acid Solution, and 20 mM HEPES (dubbed “D20”). 293T-ACE2/APN-Puro cells^39^ were cultured in D20, supplemented with 3 µg/mL puromycin. All aforementioned monolayer cell lines were cultured at 37 °C and 5% CO_2_ and passaged to maintain a density of 1-1.3 x 10^6^ cells/mL. Expi293F cells (#A14527, ThermoFisher, Waltham, MA, USA) were maintained in Expi293 expression medium at 37 °C, 8% CO_2_, and 70% humidity, shaking at 200 rpm. Expi293F cells were passaged to maintain a density of 2-3 x 10^5^ viable cells/mL. All cell lines, except Expi293F, were routinely tested for mycoplasma and remained negative.

#### Convalescent human sera and PBMCs

Sera and PBMC samples from individuals with confirmed EhCoV and SARS-CoV-2 infections were obtained under National Institutes of Health Vaccine Research Center (NIH VRC) clinical protocols, VRC200, VRC500 and VRC 317 (ClinicalTrials.gov: NCT00067054, NCT01375530, and NCT03049488 respectively). The clinical study protocols were reviewed and approved by the NIH and NIAID IRB. Written informed consent was obtained from all participants before enrollment. The studies provided compensation for participants’ time and inconvenience. Prior to conducting ELISA experiments, aliquots of frozen sera were heat-inactivated at 56 °C for 1 hr. ELISAs were performed in at least two technical replicates. Cryopreserved PBMCs were thawed in a 37 °C water bath and immediately processed in ice-cold staining buffer.

#### Mice

Animal studies were carried out in accordance with guidelines, regulations and policies for care and use of laboratory animals of the National Institute of Health. Protocols were approved by Harvard University and Harvard T.H Chan School of Public Health’s Institutional Animal Care and Use Committee. Six-to-eight weeks old female BALB/cJ mice (#000651, Jackson Laboratory, Bar Harbor, ME, USA) were housed in groups of five under a photoperiod (12 h on:12 h off dark/light cycle), ambient RT of 18 – 22 °C, and room humidity of 50 ± 5%. Mice were fed standard chow diets. Blood sampling was performed via terminal cardiac puncture under anesthesia, and efforts were made to minimize animal suffering.

## METHOD DETAILS

### Gene synthesis and plasmid construction

Chimeric EhCoV IDDs were constructed by GenScript (Piscataway, NJ, USA). IDDs were genetically fused to the N-terminus of the trimeric I53-50A.1NT1 NP component^29^ using a linker (GGGGSAEAAAKASSAEAAAKEAAAKEAAAKEEAARK). The IDD-I53-50A.1NT1 NP constructs were synthesized and cloned by GenScript (Piscataway, NJ, USA) into a pcDNA 3.1(+) mammalian expression vector for protein production in Expi293F cells. All IDD-bearing components contained an N-terminal signal peptide and a C-terminal streptag II and octa-histidine tag.

For antigenicity and competition assays: plasmid encoding hACE2-Fc^92^ was provided by Jason McLellan (University of Texas at Austin). Plasmids encoding heavy and light chains of mAbs (H501-003, −007, −008, −009, −010, −012, −013, −014, −015, −016, −018, −020, −022, and −101) (Wang, *et. al*., submitted) were synthesized in-house. Plasmids encoding heavy and light chains of mAb SD1.040^47^ was provided by David Robbiani (Institute for Research in Biomedicine). Plasmids expressing heavy and light chains of mAbs, SA58^73^, S309^70^, VIR-7229^68^, D12^28^, and F07^28^, were synthesized and cloned into pcDNA 3.1(+) mammalian expression vector by GenScript (Piscataway, NJ, USA).

For B-cell immunodominance and domain immunogenicity studies, DNA fragments encoding NTD and RBD from HKU1-CoV S protein gene (GenBank accession, ABC70719.1) were cloned into pcDNA 3.1(+) mammalian expression vector by GenScript (Piscataway, NJ, USA). Both constructs were modified to incorporate an N-terminus signal peptide (MDAMKRGLCCVLLLCGAVFVSA), a C-terminus Avi-tag (GLNDIFEAQKIEWHE) for site-directed biotinylation, and octa-histidine tag (HHHHHHHH) for purification. A human rhinovirus 3C protease recognition site (GSRSLEVLFQGP) was inserted to enable tag cleavage after purification in preparation for animal immunization.

For ELISAs, multiplex antigen binding, and Fc-effector function assays, we expressed proteins using previously reported^93^ codon-optimized contemporary S protein sequences from SARS-CoV (GenBank accession, AAP13441.1), SARS-CoV-2 (GenBank accession, QHD43416.1), WIV1-CoV (GenBank accession, KC881007), HKU1-CoV (GenBank accession, ABC70719.1), OC43-CoV (GenBank accession, AIL49484.1), 229E-CoV (GenBank accession, NP073551.1), NL63-CoV (GenBank accession, YP 003767.1), and MERS-CoV (GenBank accession, AFY13307.1); plasmids were synthesized in-house. For clade-specific quantification, selected codon-optimized S protein sequences for clades A and B (GenBank accession, **Supplementary Fig. 2**) of SARS-CoV-2 (residues 1-1214; 1-1206), HKU1-CoV (residues 1-1176;1-1176), OC43-CoV (residues 1-1293;1-1287), 229E-CoV (residues 1-1115;1-1115) and NL63-CoV (residues 1-1291;1-1291) were synthesized and cloned into pcDNA 3.1 (+) vector by GenScript (Piscataway, NJ, USA). All S proteins constructs contain a C-terminal foldon trimerization motif (YIPEAPRDGQAYVRKDGEWVLLSTFL) and an octa-histidine tag (HHHHHHHH) for purification. A human rhinovirus 3C protease recognition site (GSRSLEVLFQGP) was inserted to enable tag cleavage after purification. In addition, S proteins were prefusion stabilized via two proline substitutions and contain S1/S2 furin cleavage site modification^25,26^.

For lentivirus or vesicular stomatitis virus (VSV) pseudoviruses, codon-optimized full-length S plasmids for SARS-CoV-2 Beta-B.1.351(GenBank accession, QRN78347.1), Delta-B.1.617.2(GenBank accession, QUD52764.1), BA.5(GenBank accession, UOZ45804.1) BQ.1.1(GenBank accession, UWM38596.1), XBB.1 (GenBank accession, WIL50295.1), and BA.2(GenBank accession, ULB15050.1), HKU1-CoV (GenBank accession, DQ339101), WIV1-CoV (GenBank accession, KC881007), 229E-CoV (GenBank accession, NP_073551.1), and NL63-CoV (GenBank accession, NC_005831) were cloned into pCMV vector. These S protein pCMV vectors, pHR’ CMV Luc (luciferase reporter gene), and pCMV DR8.2 (lentivirus backbone) were constructed as previously described^25^. S protein plasmids for SARS-CoV-2 KP.3 (GenBank accession, XDZ73367.1) EG5.1(GenBank accession, WGM84363.1) and XBB.1.5(GenBank accession, UZG29433.1) were constructed as previously described^94^ and obtained from Alejandro Balazs (Ragon Institute). Codon-optimized S protein plasmids for SARS-CoV-2 Wuhan-Hu-1(GenBank accession, QHD43416.1), and JN.1 (GenBank accession, XBA04983.1) were synthesized containing 19 amino acid c-terminal deletion and cloned into pcDNA 3.1(+) vector by GenScript (Piscataway, NJ, USA). For clade-specific lentiviral or VSV plasmids, codon-optimized S plasmids for HKU1-CoV (GenBank accession, ARB07608.1), NL63-CoV (GenBank accession, QEG03748.1), and 229E-CoV (GenBank accession, QEG03785.1), containing respective 17-,18- and 16-amino acid C-terminal deletions, were synthesized and cloned into pcDNA 3.1(+) vector by GenScript (Piscataway, NJ, USA). These C-terminal deletions are known to improve viral production and infectivity^95^.

### Bioinformatics and computational analysis

For phylogenetic analysis, sequences of full-length S were derived from nextstrain (https://nextstrain.org/) and filtered by time frame. Next, sequences were aligned by Clustal Omega^96^ to develop a multiple sequence alignment (MSA). Phylogenetic trees were constructed with Clustal Omega EMBL-EBI (https://www.ebi.ac.uk/jdispatcher/msa/clustalo) with default settings using a seeded guide trees and HMM profile-profile technique. EhCoVs clades were determined by divergence timeline and evolution of major lineages. Candidate spikes from both clades were selected based on their unique position on the tree. Candidate spikes centrally positioned were hypothesized to have better cross-reactive potential. For evolutionary conservation calculation and profiling in **Fig. 4C**, ConSurf server was used (https://consurfdb.tau.ac.il/). SARS-CoV-2 RBD-SD1 structure (PDB:8D48) was uploaded, and the resultant molecular structure was colored based on sequence conservation as determined by ConSurf^97^.

### Antibody footprint, structural analysis and comparison

All protein structures used in this work were retrieved from the RCSB protein databank (PDB) (https://www.rcsb.org/).The SARS-CoV-2 S (PDB:6VXX) and RBD-SD1 domain (PDB:8D48) structures were used to showcase mutations within variants, epitope footprints, and antibody comparisons in **Fig. 4D**.

To analyze and compare antibody epitopes, RBD or SD1 domains within the antibody complex structures of CC12.1 (PDB:6XC2), B38 (PDB:7BZ5), CB6 (PDB:7C01), C102 (PDB:7K8M), C002 (PDB:7K8S), LY-CoV 555 (PBD:7KMG), P2B-2F6 (PDB:7BWJ), 47D11 (PDB:7KAJ), n3113 (PDB:7VNB), SA58 (PDB:7Y0W), S309 (PDB:6WS6), CR3022 (PDB:6W41), COVA1-16 (PDB:7JMW), VIR-7229 (PDB:9ASD), SD1.040 (PDB:8D48), P008-60 (PDB:7ZBU), and SD1-3 (PDB:8R1D) were used. Structural figures were generated using UCSF ChimeraX^98^ and epitopes were identified using default parameters. ePISA (http://www.ebi.ac.uk/pdbe/prot_int/pistart.html) was used to identify interface residues.

### Spike and immunogen expression and purification

S protein ectodomains, HKU1-CoV RBD and NTD probes, and all IDD-153-50 A.1NT1 components were produced as previously described with minor modifications^25,93^. Briefly, Expi293F cells, grown to a density of 3 x 10^6^ cells/mL, were transfected using 1 mg/mL of polyethyleneimine “PEI-MAX” (#24765, Polysciences, Warrington, PA, USA) and DNA plasmids in a 3:1 (w:w) ratio. Cells were cultured at 37 °C, 8% CO_2_, 80% humidity, and shaking at 120 rpm in polycarbonate baffled flasks. For SARS-CoV-2 IDD-L-I53-50 A.1NT1, protein was expressed at 34°C under similar conditions. Next, supernatants were collected five days post-transfection and clarified via centrifugation at 4000 xg for 30 mins. Clarified supernatants were filtered with 0.22-µm filter (#4612, Pall Corporation, Port Washington, NY, USA), diluted, and incubated at 4°C overnight with Ni-NTA agarose beads (# 30210, Qiagen, Germantown, MD, USA) for purification. Resins were washed with 10 column volumes of wash buffer (1X PBS with 20 mM imidazole). Proteins were eluted with elution buffer (1X PBS with 300 mM Imidazole) and concentrated in 10-KDa cutoff centrifugal device (#MAP010C37, Pall Corporation, Port Washington, NY, USA). Subsequently, all proteins were further purified via size-exclusion chromatography (SEC) using Superdex 200 Increase (#28990944, Cytiva, Marlborough, MA, USA) or Superose 6 increase 10/300 (#29091596, Cytiva, Marlborough, MA, USA) columns on an AKTA pure (Cytiva, Marlborough, MA, USA) in running buffer (1X PBS with 5% glycerol). Proteins were flash-frozen in liquid nitrogen and stored at −80°C until further use. Protein purity was assessed by SDS-PAGE. Before mouse immunizations, purification tags were removed using HRV-3C protease (#Z03092-100, GenScript, Piscataway, NJ, USA) following manufacturer’s protocol. Protein concentrations were determined with Nanodrop Lite spectrophotometer (ThermoFisher, Waltham, MA, USA) using the protein’s peptidic molecular weight and extinction coefficient, obtained from online Expasy ProtParam software (https://web.expasy.org/protparam/). For B-cell experiments, HKU1-CoV NTD and RBD proteins were further biotinylated using EZ-Link™ Sulfo-NHS-LC-Biotinylation Kit (#21435, ThermoFisher, Waltham, MA, USA) following manufacturer’s protocol. Biotinylated proteins were further purified via SEC, flash-frozen, and stored in aliquots at −80 °C until further use.

### Antibody expression and purification

Monoclonal antibodies (mAbs) were produced by transiently co-transfecting Expi293F cells with heavy and light chain plasmids using polyethyleneimine “PEI-MAX” (#24765, Polysciences, Warrington, PA, USA). Six days post-transfection, Expi293F supernatants were harvested, centrifuged, and filtered using 0.22-µm filters. mAbs were purified with MabSelect PrismA agarose beads (#17549801, Cytiva, Marlborough, MA, USA) using standard procedures^93^. Bound mAbs were eluted with 100 mM glycine at pH 2.7 into 1/10th volume of 1 M Tris-HCl pH 8.0, concentrated, and buffer exchanged using 10-KDa cutoff centrifugal devices into 1X PBS, pH 7.4. Concentrations were determined using Nanodrop Lite spectrophotometer (ThermoFisher, Waltham, MA, USA). Molecular weights and extinction coefficients were calculated using the online ExPASy ProtParam tool (https://web.expasy.org/protparam/). mAbs were validated by ELISA binding to apt proteins before they were flash-frozen in liquid nitrogen and stored at - 80 °C.

### I53-50B.4PT1 expression and purification

I53-50B.4PT1 was produced essentially as previously described^29^. Briefly, Lemo21 (DE3) (#C2528J, NEB, Ipswich, MA, USA) cells transformed and expressing I53-50B.4PT1 were cultured in a 2L shaker flask. Cells were cultured in LB media (10 g Tryptone, 5 g Yeast Extract, 10 g NaCl) at 37 °C to an OD_600_ of 0.8-1. Next, temperature was reduced to 18 °C, protein expression was induced with 1 mM IPTG, and cells were cultured for 20 hrs. Culture was collected and harvested via centrifugation at 10,000 xg. The resulting pellet was lysed in a lysis buffer (50 mM Tris, 500 mM NaCl, 30 mM imidazole, 1 mM PMSF, 0.75% CHAPS) via ice sonication (amplitude - 50%, time - 5 mins, pulse duration – 15 secs ON + 45 secs OFF). Lysate was clarified by centrifugation at 20,000 xg for 20 mins, and clarified supernatants were incubated with Ni-NTA agarose beads for 1 hr. Protein-bound Ni-NTA agarose beads were washed with ten column volumes of 1X PBS, 0.75% CHAPS buffer, and protein was eluted with elution buffer (50 mM Tris, 500 mM NaCl, 300 mM imidazole, 0.75% CHAPS). Eluted protein was immediately concentrated with a 10-KDa cutoff centrifugal device, filtered with a 0.22-µm filter, and further purified via SEC on Superdex 200 Increase 10/300 (#28990944, Cytiva, Marlborough, MA, USA). Purified protein was concentrated, aliquoted, flash-frozen, and stored at −80 °C until further use. Endotoxin levels were confirmed to be less than 50 EU/mL prior to NP assembly.

### *In vitro* assembly and purification of nanoparticle immunogens

NPs were made as previously described^54,99^. To assemble single EhCoV IDD NPs, IDD-I53-50A.1NT1 and I53-50B.4PT1 components were mixed at a 1.1:1 molar ratio and incubated for 1 hr at RT. To assemble multivalent mosaic NPs, three or five IDD-I53-50A.1NT1s and I53-50B.4PT1s were combined at an equimolar ratio. Then, the mixtures were mixed with I53-50B.4PT1 at a 1.1:1 molar ratio and incubated for 1 hr at RT. Assembled mixes were filtered and applied to a Superose 6 increase 10/300 column (#29091596, Cytiva, Marlborough, MA, USA) on an AKTA pure in running buffer. Fractions corresponding to the assembled NPs were pooled, concentrated using 30-KDa centrifugal devices (#MAP030C37, Port Washington, NY, USA), flash-frozen in liquid nitrogen, and stored at −80 °C until further use. NP concentrations were determined with a Nanodrop Lite spectrophotometer (ThermoFisher, Waltham, MA, USA) using the peptidic molecular weight, extinction coefficient, and formula previously reported^100^.

### Generation of HKU1-CoV S1 domains probes

To produce tetramer probes with uniformly oriented RBD and NTD domains, purified biotinylated proteins were incubated with tetrameric fluorescently labelled streptavidin-phycoerythrin (PE) (#554061, BD Biosciences, Woburn, MA, USA) and streptavidin-PerCP-Cyanine5.5 (PerCP-Cy5.5) (#551419, BD Biosciences, Woburn, MA, USA), respectively, on ice for 1 hr at a molar ratio of 6:1. Mass ratios were calculated as previously described^101^. Decoy PE or PerCP-Cy5.5 probes were prepared via conjugation to irrelevant biotinylated Ovalbumin (Ova) protein with streptavidin PE or PerCP-Cy5.5. The mixtures were purified on Superose 6 increase 10/300 column (#29091596, Cytiva, Marlborough, MA, USA) on an AKTA pure to remove excess or unconjugated streptavidin and biotinylated proteins. Tetramer fractions were pooled and concentrated in 100-KDa cutoff centrifugal devices (#MAP100C37, Pall Corporation, Port Washington, NY, USA). Tetramer concentrations were determined by measuring absorbance of PE and PerCP-Cy5.5 at 565 nM and 695 nm, respectively. Tetramers were made fresh for each experiment.

### Single B-cell flow cytometry

Flow cytometry analysis of B-cells was performed as previously described^101^ with minor modifications. Briefly, B-cells from convalescent human PBMCs were enriched using the pan-B-cell isolation kit (#130-101-638, Miltenyi Biotec, Charlestown, MA, USA) following manufacturer’s recommendations. The enriched B-cells were resuspended in fluorescence-activated cell staining FACS buffer (1X PBS, 2% BSA, 1 mM EDTA) containing Fc blocking reagent (#130-059-901, Miltenyi Biotec, Charlestown, MA, USA) and live/dead eBioscience fixable viability dye eFluor 780 (#65-0865-18, Thermo Fisher Scientific, Waltham, MA, USA) at a 1:300 dilutions on ice for 15 mins. Amine-reactive blue cell dye excluded dead cells, and FcR receptor prevented non-specific Fc receptor binding. Subsequently, B-cells were washed and stained with HKU1-CoV NTD or RBD probes at 1:200 dilution on ice for 1 hr. Decoy probes were added at the same dilution to distinguish antigen-specific B-cells from those binding to streptavidin. Finally, B-cells were rewashed and stained with the following antibodies/reagents/surface markers on ice for 45 mins at 1:300 dilution in FACS buffer: anti-IgD-BV421 (#348225, BioLegend, San Diego, CA, USA), anti-CD138-BV-510 (#356517, BioLegend, San Diego, CA, USA), anti-IgM-BV-605 (#314523, BioLegend, San Diego, CA, USA), anti-CD45-BUV-395 (#563791, BD Biosciences, Woburn, MA, USA), anti-IgG-BUV-805 (#742041, BD Biosciences, Woburn, MA, USA), anti-CD27-PE-CF594 (#567671, BioLegend, San Diego, CA, USA), anti-CD80-Cy7 (#305217, BioLegend, San Diego, CA, USA), anti-CD38-AF700 (#356623, BioLegend, San Diego, CA, USA), and anti-CD20-APC (#302309, BioLegend, San Diego, CA, USA). Cells were washed, fixed in 4% paraformaldehyde (PFA), rewashed, run on a FACSAria II (BD Biosciences, Woburn, MA, USA). Data were analyzed using FlowJo software v10.10 (FlowJo, LLC, Ashland, Oregon, USA).

### SDS-PAGE analysis

Protein purity and molecular weights were analyzed on SDS-PAGE. Briefly, 40 µL of purified proteins were mixed with 10 µL ß-mercaptoethanol-supplemented Laemmli loading dye (x1) (#1610747 Bio-Rad, Waltham, MA, USA), and sample mixtures were heated at 95°C for 10 mins. Next, 20 µL of the protein-dye mixtures and 5 µL of PageRuler Plus Prestained Protein Ladder (#26619, Thermo Fisher Scientific, Waltham, MA, USA) were run on SurePAGE, Bis-Tris, 4-20% gel (#M00656, GenScript, Piscataway, NJ, USA) in MOPS buffer (#M00138, GenScript, Piscataway, NJ, USA) for 1 hr at 130 V. Gels were stained in Coomassie blue, destained in deionized water, and imaged with UV ChemiDoc MP Touch Imaging System (Bio-Rad, Waltham, MA, USA).

### Dynamic light scattering (DLS)

Dynamic light scattering analysis was performed using Wyatt DynaPro Plate Reader (Wyatt Technology Corp, DynaPro Plate Reader III) at 25 °C in a 96-well Sensoplate (#655892, Greiner Bio-One, Monroe, NC, USA) in triplicate to measure hydrodynamic diameter (D_h_) and polydispersity of the IDD proteins and NPs in solution. Proteins were diluted to 500 µg/mL, filtered and loaded into wells for 10 acquisitions. Data was analyzed using instrument software.

### Differential scattering fluorimetry

#### Conventional DSF

Real-time PCR system, QuantStudio 6/7 Flex (ThermoFisher, Waltham, MA, USA) was used to monitor protein thermal shift by measuring changes in fluorescence of a dye that binds to protein as it unfolds. Briefly, all IDD trimers and NPs at 250 µg/mL and containing 5X SYPRO Orange Protein Gel Stain (#S6650, ThermoFisher, Waltham, MA, USA) were added to a MicroAmp FAST Optical 96-Well plate(#4346907, ThermoFisher, Waltham, MA, USA) in duplicate and sealed with MicroAmp optical adhesive film (#4360954, ThermoFisher, Waltham, MA, USA). Continuous fluorescence measurements at excitation and emission wavelengths (465 nm; 580 nm) were acquired over a linear temperature range of 25°C to 95°C at a ramp rate of 1°C /min. Data were fitted using instrument software and plotted as the first derivative of the melting curves. The melting temperatures were determined at the peaks of the first derivative.

#### NanoDSF

Prometheus NT.Plex instrument (NanoTemper Technologies, Watertown, MA, USA) was used to monitor intrinsic tryptophan and tyrosine fluorescence intensity at 350 nm and 330 nm as a function of temperature. Diluted IDD trimer and NPs at 250 µg/mL were loaded into Prometheus NT.Plex nanoDSF Grade High Sensitivity Capillary Chips (in duplicate) and left to melt at a ramp rate of 1°C/min over a temperature range of 25°C to 95°C. The intrinsic fluorescence and scattering signals were acquired using instrument software (NanoTemper) and data were analyzed with a software package (PR.ThermControl). Melting temperatures were determined based on peaks observed in the first derivatives of the F350/F330 ratio.

### Negative stain electron microscopy (NS-TEM)

To image NPs by negative-stain electron microscopy, bare and IDD NPs were diluted to 75 µg/mL in 1X PBS, pH 7.4. 4 µL of samples were applied to freshly glow-discharged carbon-coated 400-mesh copper grids (#FCF400-Cu-50, Electron Microscopy Sciences, Hatfield, PA, USA) and incubated for 1 min. The grids were blotted, dipped into distilled water droplets, and blotted with Whatman no.1 filter paper. Next, grids were stained with 10µL of 0.75% (w/v) uranyl formate stain for 30 sec and immediately blotted off with filter paper to remove excess stain. Finally, grids were allowed to air-dry for 1 min and imaged using Phillips CM10 Transmission electron microscope (ThermoFisher, Waltham, MA, USA) equipped with a tungsten filament and Gatan UltraScan (2k x 2k) CCD camera operating at 100 kV. Grids were imaged at 52,000x magnification.

### Cryo- electron microscopy (Cryo-EM)

#### Cryo-EM data acquisition

To obtain a cryo-EM particle reconstruction of the Mosaic-5 IDD NP, 4 µL of purified NP at 3 mg/mL in 1X PBS, pH 7.4 was applied onto a freshly glow-discharged (at 30 secs at 20 mA) Quantifoil R 1.2/1.3 Cu 300-mesh grid (#Q350CR1.3, Electron Microscopy Sciences, Hatfied, PA, USA). The grids were blotted for 4 secs at 4°C and 100% humidity using a blot force of 1 and a blot time of 8 secs. The blotted grids were immediately plunge-frozen in cryocooled liquid ethane using a Vitrobot Mark IV (ThermoFisher, Waltham, MA, USA) and stored in liquid nitrogen until imaging. The grids were imaged at 1.1 Å per pixel using Talos Arctica cryo-electron microscope (Thermo Fisher Scientific, Waltham, MA, USA) operated at 200 kV. Micrographs were recorded with a post-GIF Gatan K3 electron detector using SerialEM version 4.1. Each movie comprises 47 frames with an exposure time of 4.2 secs and 60.04 electron dose. A dataset of 346 movie stack or micrographs were collected in a single session with defocus range of 0.1 −2.4 µm.

#### Single-particle image processing and 3D reconstruction

All collected datasets were processed using cryoSPARC v4.6.2 package^102^ as follows. Motion correction was completed in patch motion correction module and contrast transfer function (CTF) was evaluated in patch CTF estimation. Next, ∼ 510 particles were manually picked from 50 micrographs in Manual Picker for initial 2D classification. Five 2D classes were determined and used as templates in the Template picker for automatic picking of ∼60,000 particles from 347 micrographs. Following 3 rounds of iterative 2D classification, 16,619 particles were selected and used for Ab-initio 3D reconstruction (using C1 symmetry) and heterogenous refinement enforced with or without Icosahedral symmetry. Both symmetries yielded the same cryo-EM map with well-resolved I53-50 core, but IDD trimers were not visible. In **Fig. 2E**, the sharpened cryo-EM map was visualized in UCSF ChimeraX^98^ and fitted with the computational designed model (PDB:7SGE) to enable visualization of the I53-50 scaffold and determine if displayed IDD antigens did not distort the I53-50 scaffold. All details for cryo-EM data collection, refinement, and validation statistics are available in **Supplementary Table 2**.

### Bio-layer interferometry for antigenicity

Antigenicity characterization of IDD trimers and NPs was performed and analyzed using an Octet RH96 system (Sartorius, Cambridge, MA, USA) at 30°C, shaking at 1,000 r.p.m. mAbs were diluted to 40-50 µg/mL in 1X HBS-EP+ kinetic buffer (#BR100669, Cytiva, Marlborough, MA, USA). IDD trimers were diluted to 100 µg/mL in kinetic buffer and serially diluted two-fold for a final concentration of 6.25 µg/mL. Reagents were applied to a solid black, tilted-bottom 384-well plate (#18-5076, Sartorius, Cambridge, MA, USA) at 100 µL/well. Following manufacturer’s instructions, mAbs were immobilized onto protein A biosensors, ProA (#18-5010, Sartorius, Cambridge, MA, USA), for 180 secs following initial 10 min equilibration. Next, immobilized-Ab biosensor tips were then transferred to kinetic buffer for 60 secs to reach a baseline and remove excess mAbs before subsequent binding assessments of IDD trimers via association and dissociation steps. The association step was carried out by dipping immobilized biosensors into the IDD immunogens for 300 secs, and the subsequent dissociation step was performed by dipping biosensors back into kinetic buffer for 180 secs. Data analysis and curve fitting were done with the Octet software, v8.0 (Sartorius, Cambridge, MA, USA). Data were baseline-subtracted and fitted before exporting. Experimental data were fitted with 1:1 binding model in the Octet software, v8.0 to calculate kinetic parameters. All experiments were independently performed twice, and values shown are representative means.

### Bio-layer interferometry for serum competition

Serum antibody competition was performed using an Octet RH96 instrument (Sartorius, Cambridge, MA, USA) as previously described^54,63^. The inhibition was carried out under similar conditions described above, following a detailed workflow (**Extended Data Fig. 6A**) to map and quantify Ab epitopes in sera. Pooled sera were obtained by combining equal volumes of serum from each mouse within the same vaccine group. Briefly, SARS-CoV-2, OC43-CoV, and 229E-CoV IDD trimers diluted in kinetic buffer at 20 µg/mL were loaded onto equilibrated anti-Penta-HIS biosensors, Octet HIS1K (#18-5120, Sartorius, Cambridge, MA, USA) for 5 mins to reach a capture level between 0.9 −1.2 nm. Immobilized biosensors were then equilibrated for 1 min in kinetic buffer to remove excess IDD trimers and then dipped into immune mouse sera or kinetics buffer (control) for 5 mins. Sera were diluted (starting at 1:10, 4-fold, x8) in kinetic buffer. Following this, equilibrated biosensors were dipped into 50 µg/mL competing protein or mAbs for 180 sec. Raw data for antibody competitive binding signals were analyzed, fitted with 1:1 binding model in the Octet software, v8.0 (Sartorius, Cambridge, MA, USA), exported, and plotted in GraphPad Prism v10.3.0. A representative is shown in **Extended Data Fig. 6B-C**. Ro represents the response of maximum noncompeting binding curve. Rc represents the response of the serum competing binding curve. Percent inhibition or competition was determined using the equation ([Ro-Rc]x100/Ro). All experiments were independently performed twice, and data shown are representative means.

### Immunogen-adjuvant preparation

Immunogen-adjuvant formulations were prepared as previously described^25^ with minor modifications. Purified immunogens (containing low endotoxin, < 50 EU/mL) concentrations were determined again with a Nanodrop using the peptidic molecular weight and extinction coefficient prior to formulation. Each mouse immunized with IDD or IDD NPs received 5µg IDD or corresponding molar equivalent of IDD-NP with equal IDD weights. An oil-in-water emulsion adjuvant (SAS #S6322, Sigma, Burlington, MA, USA) was used. Prior to formulation, SAS adjuvant was warmed at 42 °C and prepared in sterile 1X PBS following manufacturer’s recommendations. Next, immunogens diluted in 1X PBS and combined with SAS adjuvant at a volume ratio of 1:1; the PBS group was also mixed with SAS. Formulated immunogen-adjuvant combos were briefly vortexed at low speed, thoroughly mixed on a shaker for 20 mins and stored at 4 °C shaking for at least 30 mins prior to immunization.

### Endotoxin removal and measurements

Endotoxin removal and testing were completed as previously described^54^. Briefly, endotoxin was removed from I53-50B.4PT1 component during protein purification by washing with detergent-containing buffer (1X PBS, 0.75% CHAPS) during Ni-NTA affinity chromatography. Purified I53-50B.4PT1 pentamer was tested for endotoxin prior to NP formulation using Kinetic-QCL™ Kinetic Chromogenic Limulus Amebocyte Lysate (LAL) kit (# 50-650U, Lonza Bioscience, Cambridge, MA, USA) following manufacturer’s recommendations. Prior to and following NP assembly and formulation, proteins were again tested for endotoxin contamination. The endotoxin concentrations measured were negative or below < 50 EU/mg, a sample threshold suitable for immunization.

### Mouse immunizations

Immunizations were completed using standard timeline and protocol as previously described^93^. BALB/cJ mice, as described above, were immunized IM at weeks 0 and 3 with 50 μL into each hind leg. Mice were bled via humane endpoint terminal bleeding by cardiac puncture at week 5. Blood samples were allowed to clot for 30 mins, and sera were collected and stored at −80 °C or heat-inactivated at 56 °C for 1 hr to evaluate immune response via subsequent serological assays such as ELISA, neutralization, and BLI competition assay.

### Anti-sera enzyme-linked immunosorbent assay (ELISA)

Cross-reactive binding Ab responses were evaluated using ELISA as previously described^92^ with minor modifications. For mouse anti-sera ELISA, 96-well EIA/RIA clear flat bottom microplates (#3361, Corning, Tewksbury, MA, USA) were coated with 1 µg/mL of S protein in 100 μL in 1X PBS and incubated at 4 °C for 16 hrs. When testing for binding to CoV S, S-2P design^25^ was used. Following standard washes in PBS-T (1X PBS + 0.05% Tween 20) buffer using AquaMax microplate washer (Molecular devices, San Jose, CA, USA) and blocks in blocking buffer (1X PBS-T + 5% nonfat Difco skim milk, #232100, Midland Scientific, Vista, NE, USA) for 1 hr at RT, coated plates were incubated with serially diluted (starting at 1:100), heat-inactivated mouse sera for 1 hr at RT. Following standard washes with PBS-T buffer, 1:4000 dilution of goat anti-Mouse IgG (H+L) cross-adsorbed secondary Ab HRP (#G-21040, ThermoFisher, Waltham, MA, USA) was added to the plates and incubated at RT for 1 hr. Finally, the plates were rewashed 3x with PBS-T and developed using 100 µL of 1-Step 3,5,3′5′-tetramethylbenzidine (TMB) ELISA peroxidase substrate solution (#N301, ThermoFisher, Waltham, MA, USA) per well for 10 mins to detect Ab response. Next, the plates were quenched with 100 μL of 1N sulfuric acid (# SA212-1, ThermoFisher, Waltham, MA, USA) and read at 450 nm and 650 nm using SpectraMax iD5 Multi-Mode Microplate Reader (Molecular Devices, San Jose, CA, USA). The datasets were analyzed and exported using SoftMax Pro Software. Endpoints titers were determined as the highest dilution that showed A_450_ as 4-fold above the background average (secondary Ab only). For human anti-sera ELISA, 100 μL of 1 μg/mL SARS-CoV-2 IDD, SARS-CoV-2 IDD NP, Mosaic-3, or Mosaic-5 IDD NPs were plated onto 96-well microplates in 1X PBS and incubated at 4°C for 16 hrs. All following methods were completed as described above for the mouse anti-sera ELISA.

### Pseudovirus production, entry and serum neutralization assays

Pseudovirus neutralization assays were completed using lentivirus-based reporter viruses for all EhCoVs strains, except HKU1 where a VSVΔG-based system was used. These assays measure the inhibition of pseudovirus attachment, entry and fusion, and correlate with live virus plaque-reduction neutralization assay^103^. Moreover, it was preferentially selected for neutralization activity measurement as it does not require lucrative biosafety level 3 (BSL3) containment, for high-containment strains.

#### Lentivirus-based pseudovirus-neutralization assay

Pseudotyped lentiviral reporter viruses were produced as previously described^39,92^. Briefly, pseudoviruses bearing the contemporary and clade-specific codon-optimized spike glycoprotein of NL63-CoV, 229E-CoV, WIV1-CoV, and SARS-CoV-2 variants, and carrying a firefly luciferase (Luc) reporter gene were produced by co-transfecting 90% confluent HEK-293T/17 cells with 17.5µg of pCMVΔR8.2 (lentivirus backbone), 17.5 µg of pHR’CMVLuc (luciferase reporter), and 1 µg of spike, S gene plasmids seeded in 150 mm tissue culture dishes using using fugene 6 transfection (# E2692, Promega, Madison, WI, USA) in OPTI-Minimum Essential Media (# 31985-070, ThermoFisher, Waltham, MA, USA). For SARS-CoV-2 variants, NL63-CoV, and WIV-1-CoV pseudoviruses, 0.31µg of pCMV-TMPRSS2 (human transmembrane protease serine 2) plasmid was also co-transfected. The dishes were incubated overnight at 37 °C, 5% CO_2_ and replenished with fresh complete DMEM medium (DMEM with 10% FBS and 1% penicillin-streptomycin). Next, pseudovirus supernatants were harvested 48 hrs later, filtered through 0.45 µm low-protein binding Steriflip 0.45 μm PVDF filter (# SE1M003M00, Sigma, Burlington, MA, USA) and frozen at −80 °C. Prior to use in neutralization assay, pseudotyped viral stocks were titrated by infecting target cells for 72 hrs and luciferase activity was determined using Luciferase Assay System (# E1501, Promega, Madison, WI, USA). Pseudovirus titers were expressed as relative luminescence unit (RLU). 293T-hACE2 cell was used for SARS-CoV-2 variants, WIV1-CoV and NL63-CoV assays. 293T-ACE2-APN cell was used for 229E-CoV assay.

Pseudotyped virus neutralizing activity was tested as previously described ^92,93,104^. Target cells were seeded in 96-well black and white plates (#6005068, Revvity, Waltham, MA, USA) at 5000 - 10,000 cells/well and incubated at 37 °C and 5% CO_2_ overnight. On the day of infection, serial dilutions of sera samples (1:40, 4-fold, 8x), were prepared in 90 μL DMEM in U-bottom dilution plates (#267334,ThermoFisher, Waltham, MA, USA) and incubated with addition of 90 μL of pseudoviruses (at 100,000 RLU). The pseudovirus/sera mixture was incubated at 37 °C for 45 mins. 50 μL of pseudovirus/sera mixture was used to inoculate previously seeded target cells and incubated for 2 hrs at 37 °C, in triplicate. Next, the plates were topped up with the target media and pseudovirus infectivity was determined 72 hrs post-incubation for luciferase activity using Luciferase Assay System. Cells were then lysed using luciferase cell culture lysis 5X Reagent (#E1531, Promega, Madison, WI, USA) and luciferase substrate (#E1501, Promega, Madison, WI, USA) were added. Luciferase readout was measured in RLU at 570 nm using a SpectraMax iD5 Multi-Mode Microplate Reader (Molecular Devices). With uninfected cells representing 100% neutralization and cells transduced with pseudotyped virus alone representing 0% neutralization, averages of the triplicates were normalized, and sigmoidal curves were produced. Neutralizing Ab titers at 50% (ID50) were calculated by applying a nonlinear regression model [log(agonist) vs. normalized response, variable slope] to normalized values in GraphPad Prism v10.3.

#### Recombinant HKU1 VSVΔG-based pseudovirus neutralization assay

Pseudotyped HKU1-CoV was generated as previously described^27,105^ with minor modifications. Full-length codon-optimized spike gene of human coronavirus HKU1 A (GenBank accession, ARB07608) and HKU1 C (GenBank accession, DQ339101) plasmid constructs were used. Briefly, HEK-293T cells at 70-80% confluency were seeded and transfected with 10 μg of the plasmid construct using polyethylenimine (PEI-MAX) at a 1:3 ratio in OPTI-Minimum Essential Media. Cells were infected with VSV-G pseudotyped VSVΔG bearing the firefly (Photinus pyralis) Luc reporter gene at an MOI of 1, 24 hrs post-transfection and incubated for 2 hrs at 37 °C, 5% CO_2_. The virus inoculum was removed, and cells were washed with 1X PBS and supplemented with complete DMEM (DMEM with 10% FBS and 1% penicillin-streptomycin) with anti-VSV-G antibody (I1-mouse hybridoma supernatant diluted to 1:25, #CRL-2700, ATCC, Manassas, VA USA). Cell supernatant was harvested 24 hrs post-infection, cleared of cell debris, filtered (0.45μm), and aliquots were stored at −80 °C until used.

For sera neutralization assays, HEK293T-TMPRSS2 cells were seeded at 15,000 cells per well in 96-well black and white plates and incubated at 37 °C with 5% CO_2_ overnight. Serial dilutions of sera samples (1:40, 4-fold dilution, 8x) were prepared in 90 μL DMEM in U-bottom dilution plates, followed by the addition of 90 μL of HKU1-CoV pseudovirus (at 100, 000 RLU). The pseudovirus/sera mixture was incubated at 37 °C for 45 min and 50 μL of pseudovirus/antibody mixture was added to HEK293T-TMPRSS2 cells and incubated for 2 h at 37 °C followed by the addition of 100 μL of complete DMEM medium. Cells were lysed twenty hours post-infection using cell lysis buffer following the manufacturer’s instructions, and subsequently, 50 μL of luciferase substrate was added to the cells. Luciferase readout was measured in RLU at 570 nm using a SpectraMax iD5 Multi-Mode Microplate Reader. Data analysis is the same as described above for the lentivirus-based pseudovirus neutralization assay.

### Antibody subclassing/Isotyping and Fc-receptor binding profiling

Mouse sera were analyzed by a custom-built Luminex assay to quantify the levels of antigen-specific antibody features as previously described^77^. Target S-2P antigens were from contemporary or vaccine-mismatched SARS-CoV, SARS-CoV-2, WIV-1-CoV, HKU1-CoV S-2P, OC43-CoV S-2P, 229E-CoV, and NL63-CoV. Target IDD antigens were from clade-specific or vaccine-matched SARS-CoV-2, HKU1-CoV, OC43-CoV, 229E-CoV, and NL63-CoV. EBOV GP was used as a negative control. Antigens were covalently coupled with a distinct bead region of Magplex microspheres using EDC (#A35391, ThermoFisher, Waltham, MA, USA) and Sulfo-NHS (#A39269, ThermoFisher, Waltham, MA, USA). After antigen coupling, these bead regions were mixed for multiplexing. Prior to running all samples, a dilution titration was run to identify adequate serum dilution for each Ig isotype and Fc-receptor within the linear range of detection. The diluted serum samples were then added to antigen-coupled beads mixture to form immune complexes in 384-well plates (in duplicate) and incubated at RT for 2 hrs in Assay Buffer (0.1% BSA and 0.02% Tween 20 in 1X PBS, pH = 7.4), shaking at 750 rpm.

After this incubation, the plates were washed three times using Assay Buffer containing 0.1% BSA and 0.02% Tween 20 in PBS on the Tecan Hydrospeed Plate Washer [in order of aspirate, (soak, aspirate, dispense, soak, shake x3], aspirate. *see protocol details below*). Following washing, Goat Anti-Mouse antibody isotype, bound with PE for detection (IgG1, IgG2a, IgG3, IgM, IgA, Southern Biotech, Birmingham, AL, USA), was diluted in Assay Buffer and added to the plates and incubated for 1 hr at RT, shaking at 750 rpm. Fc receptors were added as secondary reagents, similar to the Igs. Prior to the assay, mouse Fc-receptors (FcγR2b, FcγR3, and FcγR4 - synthesized by Duke University, Durham, NC, USA) were biotinylated using the BirA500 kit (# NC2047451, Avidity, LLC, Aurora, CO, USA). Biotinylated Fc receptors were mixed, incubated with Streptavidin PE, added to the plates in Assay Buffer, and incubated for 1 hr at RT, shaking at 750 rpm. After the secondary reagent was added and incubated, another wash cycle was performed using buffer (x1 PBS, 0.1% BSA, 0.02% Tween 20) on the Tecan Hydrospeed Plate Washer (*see protocol details below*). Finally, 1X Luminex Sheath reagent was used to resuspend the well contents to prepare for flow cytometry acquisition on Luminex xMAP INTELLIFLEX DR-SE. The system software was used to determine the median fluorescence intensity (MFI), and results show average values of the duplicates. The protocol or setting used for the wash cycle for the Tecan Hydrospeed Plate Washer was in this order: Aspiration rate-4, Cycles-1, soak-30 secs, Cycles-3, Aspirate-1secs; 4.5 mm, Disperse-60 µL:rate=100 µL/secs, Soak and Shake-Medium:10secs, Soak-40secs, Cycles-1, Aspirate-1secs; 4.5 mm.

### Evaluation of antibody Fc-mediated effector functions

#### Antibody-dependent cellular phagocytosis and neutrophil phagocytosis (ADCP and ANDP)

A flow-cytometry-based phagocytic assay was used to evaluate ADCP and ADNP. Briefly, the same antigens used in antibody binding assay were separately biotinylated using the EZ-link Sulfo-NHS-LC-LC-Biotin kit (#21338, ThermoFisher, Waltham, MA, USA) cleaned up by running biotinylated antigens through a zeba column, and then conjugated to Yellow or Green Fluorescent Neutravidin Microspheres (#F8776,ThermoFisher, Waltham, MA, USA) using 0.1% BSA in 1X PBS to wash and resuspend the antigen-bead conjugation. Conjugated beads were then added to 96-well round-bottom plates, mixed with mouse sera (diluted 1:100 in 1X PBS), and incubated for 2 hrs at 37 °C to form immunecomplexes. Following incubation, plates were washed with 1X PBS, centrifuged at 200 xg for 10 mins, and unbound antibodies were removed. Isolated cells from human leukopak or whole blood (*see cell isolation protocol below*), 25,000 cells per well for ADCP and 50,000 cells per well for ADNP, were added to plates and incubated for 2 hrs at 37 °C. Once incubation was complete, the plates were centrifuged at 500 xg for 5 mins, and supernatants were removed. A cocktail containing 1:100 diluted CD14 Pacblue Antibody (#325619, Biolegend, San Diego, CA, USA) for ADCP or a cocktail containing 1:100 diluted CD66b V450 Pacblue Antibody (#305112, Biolegend, San Diego, CA, USA) for ADNP was then added to plates for staining cells and incubated at RT for 10 mins. Plates were then rewashed by centrifuging at 500 xg for 5 mins, and the supernatants were removed. The cells were then fixed in 4% paraformaldehyde (PFA) and incubated at RT for 10 mins. Following a second wash by centrifuging at 500 xg for 5 mins, the supernatants were removed, and cells were resuspended in PBS in preparation for flow cytometry acquisition. The iQue 3 HTS Cytometry Platform (Sartorius, Cambridge, MA, USA) was used to determine the fluorescence or phagoscore of gated monocytes and neutrophils. The cell isolation was performed as described below.

For Antibody Dependent Cellular Phagocytosis (ADCP), monocytes were isolated from a fresh Human Peripheral Blood Leukopak (#70500.2, Stemcell, Cambridge, MA, USA) using the Red Blood Cell (RBC) Depletion kit (#18170, Stemcell, Cambridge, MA, USA). First, EDTA (#15575-038, ThermoFisher, Waltham, MA, USA) was added to the leukopak to help preserve cell viability before diluting the sample. Then, depletion reagent was added to the blood sample and mixed thoroughly, labelling all unwanted cells with magnetic particles. Using an EasySep magnet, supernatant was collected from the sample, leaving unwanted red blood cells (RBCs) behind. This step was performed twice to make sure all RBCs are left behind in discarded tubes.

For Antibody Dependent Neutrophil Phagocytosis (ADNP), neutrophils were isolated from fresh human whole blood (#70500.2, Stemcell, Cambridge, MA, USA) using the EasySep Direct Human Neutrophil Isolation Kit (#19666, Stemcell, Cambridge, MA, USA). After transferring the blood to a 50mL conical tube, Isolation Cocktail and RapidSpheres were added to the sample, mixed thoroughly, and then incubated for 5 mins. An EasySep Buffer (#20144, Stemcell, Cambridge, MA, USA) was used to dilute sample and mixed thoroughly before being placed in the magnet for 10 mins. Following 10 mins incubation, supernatant was collected and added to a new conical tube, leaving behind the RBCs. The process was repeated with the now enriched cell suspension, to ensure all RBCs were left behind with the magnetic particles. The use of human cells to validate murine opsinophagocytosis responses has been previously validated and reported^106^.

#### Antibody-dependent complement deposition (ADCD)

ADCD assays were performed as previously described^107^. Briefly, the same antigens used in antibody binding assay were covalently coupled with a distinct bead region of Magplex microspheres using EDC (#A35391, ThermoFisher, Waltham, MA, USA) and Sulfo-NHS (#A39269, ThermoFisher, Waltham, MA, USA). After antigen coupling, these bead regions were mixed for multiplexing and added to 384-well plates containing serum samples (1:20 dilution in duplicate), then incubated at 800 rpm for 2 hrs at RT in Assay Buffer to form immune complexes. The plate was then washed using 1% BSA in PBS using the Tecan Hydrospeed plate washer. Next, plates were incubated with lyophilized guinea pig complement (#CL4051, Cedarlane, Burlington, Canada) in gelatin veronal buffer with calcium and magnesium (#G6514, Sigma, Burlington, MA, USA) for 20 mins at 37 °C, shaking at 800 rpm. The plate was rewashed using 0.1% BSA in PBS using the Tecan Hydrospeed plate washer. Deposition of C3 complement component was evaluated by adding an anti-guinea pig C3 FITC detection antibody (#SKU:0855385, MP Biomedicals, Solon, OH, USA) and incubated for 30 mins at RT, shaking at 800 rpm. The plate was rewashed using 0.1% BSA in PBS on the Tecan plate washer and then resuspended in PBS for flow cytometry. The iQue 3 HTS Cytometry Platform was used to detect median fluorescence intensity of FITC, and final values were reported as average of the duplicates.

#### Antibody-dependent natural killer cell (NK) activation (ADNKA)

ADNKA was quantified via the surface expression of CD107a (as a marker for degranulation) and MIP-1β (as a marker for NK cell activation).ELISA plates were coated with 3 µg/mL of the same antigens used in binding assays and incubated at 37 °C for 2 hrs. Following incubation, the plates were washed thrice with PBS, blocked with 5% BSA in 1XPBS and incubated for 1 hr at 37 °C. After blocking, ELISA plates were rewashed three times with 1X PBS, and then the mouse serum samples were added at 1:25 dilution in PBS and incubated overnight at 4 °C. Following this, plates were washed three times with 1X PBS. Next, the antigen-coated, ELISA plates were incubated with NK cell preparation at 37°C, 5% CO_2_ for 5 hrs. A stain containing CD56 PE-Cy7 (#557747, BD Bioscience, Woburn, MA, USA), CD16 APC-Cy7 (#557758, BD Bioscience, Woburn, MA, USA), and CD3 PacBlue (#558117, BD Bioscience, Woburn, MA, USA) were mixed and added to the wells of 96-well V-bottom plates.

After 5 hrs incubation, NK cells from the ELISA plates were transferred to the V-bottom plates, mixed with the surface stain, and incubated for 15 mins under foil at RT. The plates were then washed twice with PBS by centrifuging at 500 xg for 5 mins, and supernatants were removed. Plates were fixed with PermA (#GAS001S100, ThermoFisher, Waltham, MA, USA) for 15 mins under foil. Following incubation with PermA, plates were washed as previously and then intracellular stain containing anti-MIP-1β PE (#550078, BD Bioscience, Woburn, MA, USA) and PermB (#GAS002S100, ThermoFisher, Waltham, MA, USA) was added for 15 mins incubation at RT under foil. The cells were resuspended in PBS in preparation for flow cytometry acquisition on the iQue 3 HTS Cytometry Platform. NK cells were gated CD56+/CD16+/CD3-. NK activation was measured as the percentage of NK cells expressing CD107a, MIP-1, and IFNγ. All experiments were performed with at least two healthy donors; the results are donors’ average. The NK cell isolation or preparation was performed as described below.

Natural Killer (NK) cells were isolated from fresh Human Peripheral Blood Leukopak (#70500.2, Stemcell, Cambridge, MA, USA) using EasySep Human NK Cell Isolation Kit (#17955, Stemcell, Cambridge, MA, USA). Sample was transferred to a 50 mL conical tube, centrifuged to remove supernatant and resuspended at 5 x 10^7^ cells/mL for isolation. An isolation cocktail was added to the sample, mixed, and then incubated at RT for 5 min. Magnetic Rapidspheres were added, the sample was diluted to the appropriate volume (50 mL) and mixed. Next, the conical tube was moved immediately to the magnet for 5 mins. Supernatant was pipetted into a new 50 mL conical to collect the enriched cell suspension, and then put on the magnet for another 5 mins to ensure all unwanted cells were removed. The enriched cell suspension was centrifuged again, supernatant was removed, and cells were resuspended in R10 media, containing RPMI-1640 (#11875093, ThermoFisher, Waltham, MA, USA), 10% FBS, 5% penicillin/streptomycin, and 5% L-glutamine with IL-15 for cell growth and maintenance and incubated overnight at 37 °C, 5% CO_2_ (at 1.5 x10^6^ cells/mL). Cells incubated overnight were adjusted to a concentration of 2.5 x 10^5^ cells/mL. A cocktail containing anti-CD107a PE-Cy5 (#555802, BD Bioscience, Woburn, MA, USA), Brefeldin A (#B7651, Sigma, Burlington, MA, USA), and Golgi Stop (#554724, BD Bioscience, Woburn, MA, USA) were mixed and added to the cells. The use of human cells to validate murine opsinophagocytosis responses has been previously validated and reported^106^.

### Quantification and statistical analysis

Statistical experimental details including all quantifications, statistical analyses and tests, animal numbers (N) are available in figure legends. For human and mouse anti-sera ELISAs and neutralization assays, sera from 7 convalescent donors or 10 BALB/cJ animals were used, and experiments were performed in triplicate. Reciprocal endpoint and reciprocal ID50 titers were transformed so all values were on a log10 scale before statistical analyses. Geometric mean titers were determined. Dotted lines indicate assay limits of detection (LOD). Datasets were analyzed using one-way ANOVA followed by Turkey’s multiple comparisons. We utilized quantile-quantile (Q-Q) probability plots to compare actual and predicted binding and neutralization titers to determine if our data were normally distributed, which is necessary for Tukey’s multiple comparison tests. All data followed a normal distribution. Statistical analyses were performed using GraphPad Prism v10.3.0. Significance is indicated with asterisks defined as *P < 0.05, **P < 0.01, ***P < 0.001, ****P < 0.0001, and non-significant groups are not shown.Systems serology quantifications were done by taking the mean of technical replicates (binding assays) or biological donors (Fc-effector assays) and, where indicated, subtracting baseline values (no serum) for each feature. All data were quality controlled to ensure that replicate values did not exceed 50% of the mean; if replicates exceeded that threshold, the data was not reported. For all binding assays, serum samples whose binding features did not exceed the baseline values (no serum) + 3 standard deviations were not considered as reportable. For all Fc-effector assays, baseline values were directly subtracted from the reported value. Statistical analyses for systems serology-generated data were done through ANOVA with Tukey’s Post-Hoc Test. Primary comparisons were each vaccinated group to the naïve cohort.

## Data and code availability

All images and data were generated and analyzed by the authors. All data supporting the study’s findings and its extended, supplementary, and source data are available in the paper. Cryo-EM reconstructions will be provided upon request and deposited in the Electron Microscopy Data Bank (EMDB). Other additional information and raw data will be made available by the corresponding author upon reasonable request. This paper did not use any code during the data acquisition. Systems serology data were analyzed through a customized R-based pipeline, and all data processing and code were used in R version 6.0.

## Material and resource availability

All requests for reagents and resources should be directed to the corresponding author, Kizzmekia S. Corbett-Helaire and will be made available after completion of a material transfer agreement (MTA).

## Acknowledgements

We would like to thank Judy Stein, Monique Young, and Kaitlyn Morabito for technology transfer, administrative, and project management support, respectively. Acquisition of human sera and PBMC samples was supported by the Vaccine Research Center, an intramural division of National Institute of Allergy and Infectious Diseases, National Institutes of Health (NIAID, NIH). We also express our sincere gratitude to the Harvard Cryo-Electron Microscopy Center for Structural Biology, Harvard Medical School Molecular Electron Microscopy Suite (MEMS), the Center for Macromolecular Interactions (CMI), and Boston Children’s Hospital FICR-PCMM Flow and Imaging Cytometry Resource for provision of service and use of respective equipment and/or facilities. We thank all members of the Harvard Center for Comparative Medicine mouse facility at Harvard T.H. Chan School of Public Health for animal husbandry assistance. Support for systems serology assays and analysis was provided by the P01AI65072 and U19AI135995 to RPM funding. This work was supported, in part, by Howard Hughes Medical Institute Freeman Hrabowski Scholars grant (to KSC), Chan Zuckerberg Initiative Science Diversity Leadership grant (No. 2022-310965 to KSC), Hans Sigrist Foundation Prize (to KSC), in-kind gifts of lab equipment, consumables, and supplies from Corning, Inc. (to KSC), and the Melvin J. and Geraldine L. Glimcher Assistant Professorship and start-up funds from Harvard T.H. Chan School of Public Health (to KSC).

## Author Contributions

CKOD and KSC conceptualized the study. CKOD designed and engineered IDDs and NPs. CKOD, AD and SM performed protein and NP production, purification and SEC analysis. CKOD and AD performed BLI antigenicity assays. CKOD performed NS-TEM, DLS, DSF, and Cryo-EM analysis. CKOD performed B-cell immunodominance flow cytometry experiments. CKOD, SM, and KSC designed animal immunization experiments; animal experiments were completed by CKOD and SM. CKOD, SM and AT performed ELISAs. CKOD and SM performed lentiviral pseudovirus neutralization assays. CKOD and JJ performed HKU1-VSVΔG pseudovirus neutralization assays. CKOD, SM, and VF performed pseudovirus neutralization data processing. CKOD and TK performed serum competition BLI assays. LRM performed system serology analysis under the supervision of RPM. CKOD, AD and AT performed computational and bioinformatics analysis. LN and IG coordinated and acquired human sera and PBMCs samples, spearheaded by LAH and LKD. KSC supervised and administrated the work. CKOD outlined and wrote the original manuscript. CKOD and KSC revised and polished the manuscript with inputs from all coauthors.

