## Extended Data for "Rational Design of Multiclade Coronavirus Spike Immunodominant Domain Nanoparticles to Elicit Broad Antibody Responses"

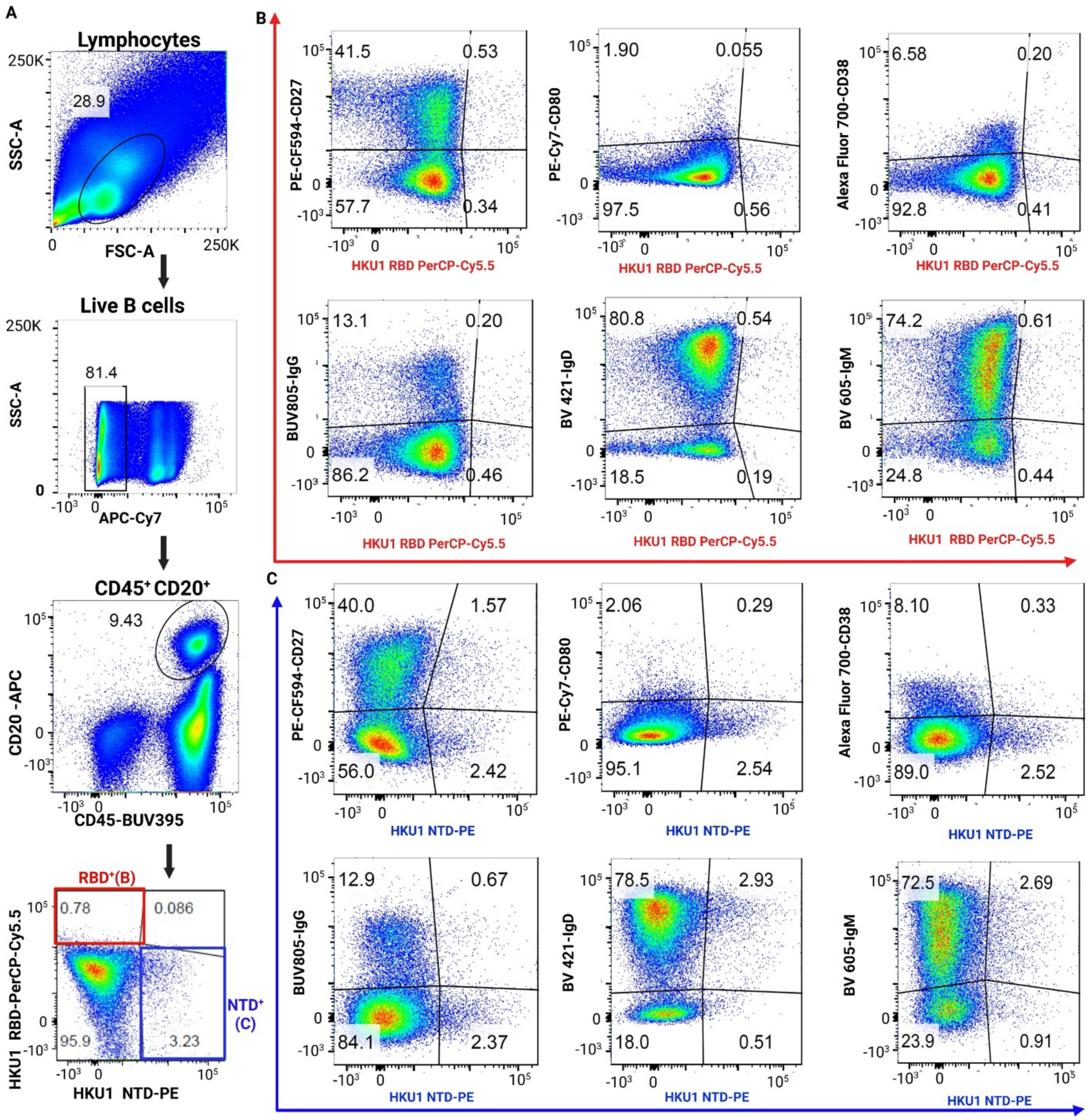


**Extended Data Fig. 1: Representative gating strategy for evaluation of HKU1-CoV S1 domain-specific B-cells by flow cytometry, related to Figure 1F.** (A) FACS analysis for identifying HKU1-CoV S RBD and NTD reactive B-cells (B-C) Representative flow cytometry plots of CD45 CD20 B-cells binding to fluorescently labelled (B) HKU1-CoV RBD probe(red) and (C) HKU1-CoV NTD probe(blue). Identification of B cell subsets(top) and Ig isotypes (bottom) within B cell population. Numbers in each quadrant indicate percentage of gated cells.

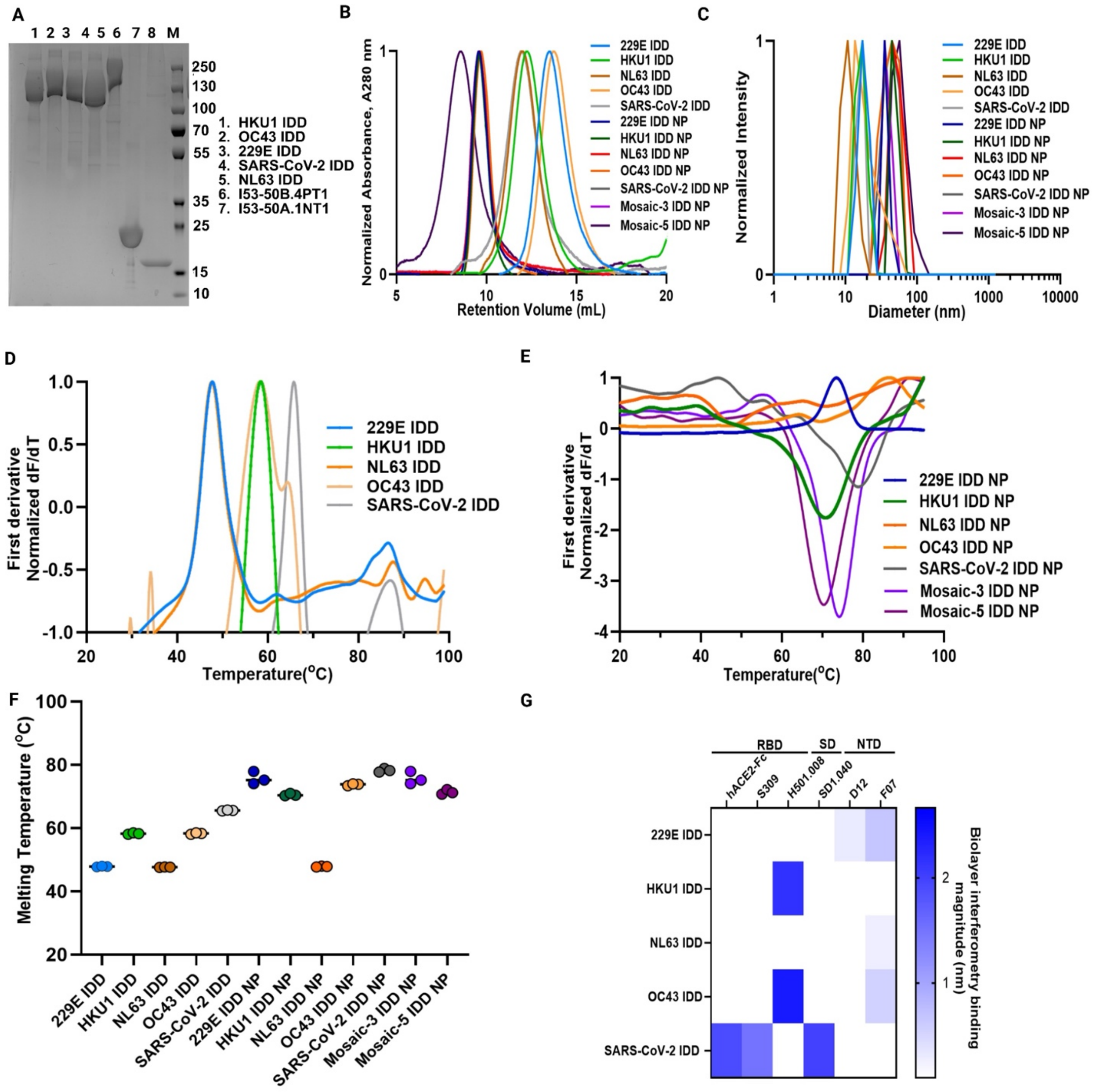


**Extended Data Fig.2: Design, production and characterization of IDD trimers and IDD NPs, related to Figs. 1H and 2.** (**A**) SDS-PAGE of affinity-purified chimeric IDD trimers, I53-50A.1NT1, and I53-50B.4PT1 subunits. (**B**) Size-exclusion chromatograms of unassembled IDD trimers and assembled NPs on a Superose 6 Increase 10/300 GL column. (**C**) Dynamic light scattering (DLS) of IDD trimers and NPs. The hydrodynamic diameter and polydispersity index (PDI) of each immunogen are shown. (**D**) Representative thermal melting profiles of IDD trimers measured by conventional differential scanning fluorimetry (DSF). Data represent the average of n=3 technical replicates. (**E**) Representative thermal melting profiles of NPs measured by nano differential scanning fluorimetry (nanoDSF). Data represent the average of three technical replicates. (**F**) Thermal melting temperatures of IDD trimers and IDD NPs measured by DSF. Data represent the average of three technical replicates. (**G**) Antigenic characterization of IDD trimers by bio-layer interferometry. Antigenicity was performed by assessing binding of immunogens to receptor, hACE2-Fc and S1 domain-specific mAbs. hACE2-Fc and S309 bind to SARS-CoV-2 RBD. H501.008 binds to HKU1-CoV and OC43-CoV RBD. SD1.040 binds to SARS-CoV-2 Subdomain, SD1. D12 and F07 bind to 229E-CoV and NL63-CoV NTD. Blue intensity signifies the magnitude of binding.

| Convalescent Sera Subject ID | | Coated Immunogens | | | | |
|---|---|---|---|---|---|---|
| | | BSA | SARS-CoV-2 IDD | SARS-CoV-2 IDD NP | Mosaic-3 IDD NP | Mosaic-5 IDD NP |
| Pre-pandemic Era | A | 2 | 2 | 2 | 3.6 | 4.6 |
| | B | 2 | 2 | 2 | 3.9 | 4.2 |
| | C | 2 | 2 | 2 | 4.7 | 4.5 |
| | D | 2 | 2 | 2 | 4.5 | 4.8 |
| COVID-19 Era | E | 2 | 2.5 | 3.8 | 3.5 | 4.2 |

| Endpoint Titer (Log10) | | | | |
|---|---|---|---|---|
| | 4 – 5 | 3 – 4 | 2.5 – 3 | 2 |

**Extended Data Fig. 3: Human convalescent sera reactivity to IDD trimers.** Sera from convalescent donors with pre-existing immunity acquired by infection in pre-pandemic and COVID-19 pandemic era were tested for IgG binding to IDD trimers. Data are presented as Logarithmic(Log10) mean endpoint titer and indicated with colors. Dark red = 4-5, red = 3-4, yellow = 2.5-3, and uncolored = 2. BSA (negative control), SARS-CoV-2 IDD trimers and IDD NPs were used to coat ELISA plates, followed by incubation with serially diluted sera (starting at 1:100) and detection with type-specific secondary antibodies. Samples were run in triplicate and performed twice.

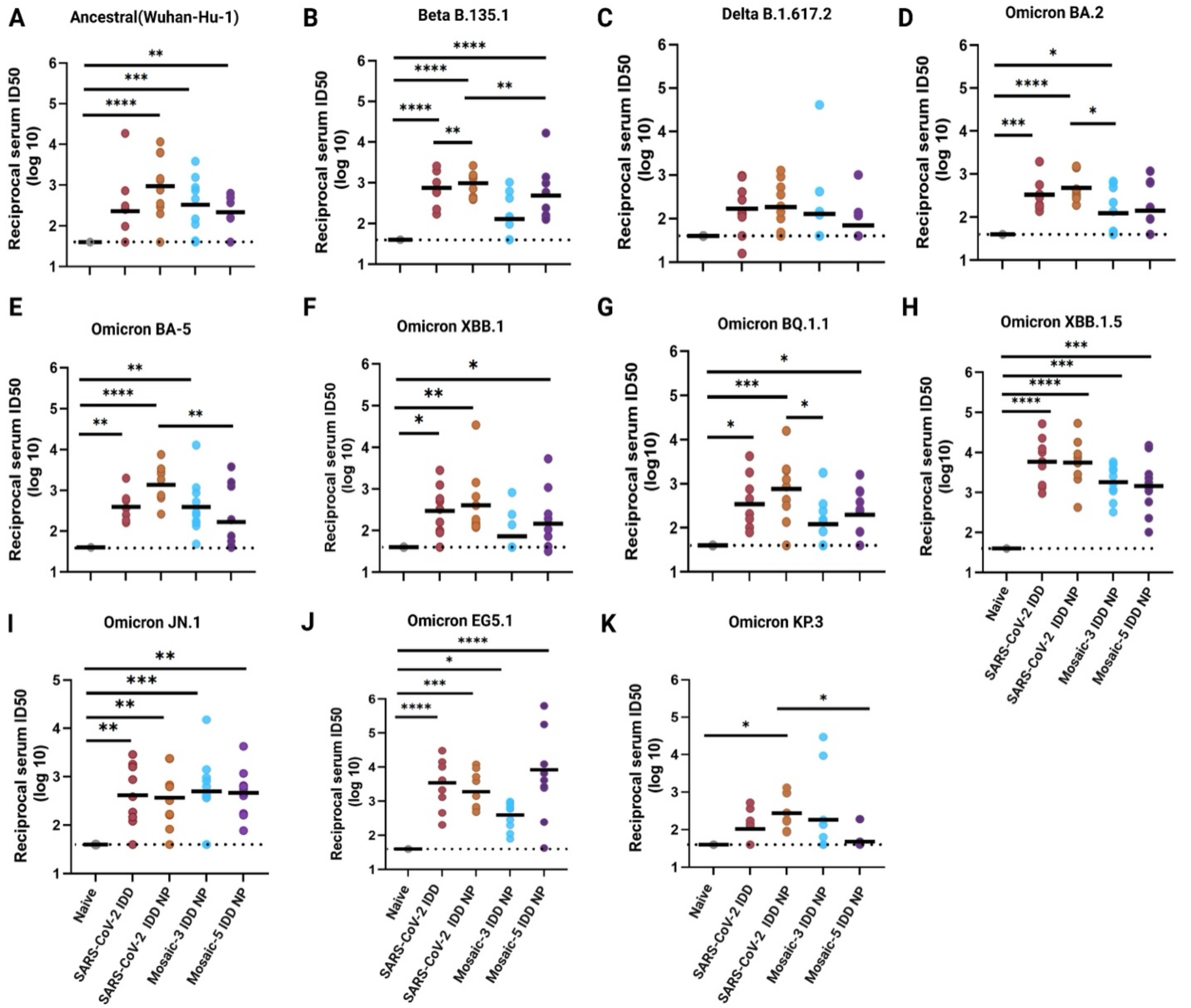


**Extended Data Fig. 4: Neutralizing antibody responses in immunized mice against SARS-CoV-2 pseudotyped viruses related to Fig. 4B, left.** Sera collected 2-weeks post immunization were evaluated for neutralization against the following pseudotyped SARS-CoV-2 strains: ancestral (Wuhan-Hu-1), Beta, Delta, Omicron BA.2, BA.5,XBB.1,BQ1.1, XBB 1.5,JN.1, EG5.1, and KP.3. Horizontal dashed lines represents the assay LOD. Bars indicates geometric mean ID50 for ten mice per group. Each dot represents an individual mouse. Two–way ANOVA with Tukey's post-hoc test was used to compare geometric mean ID50 titers. Statistical significance is displayed as follows: ∗p < 0.05; ∗∗p < 0.01; ∗∗∗p < 0.001; ∗∗∗∗p < 0.0001.

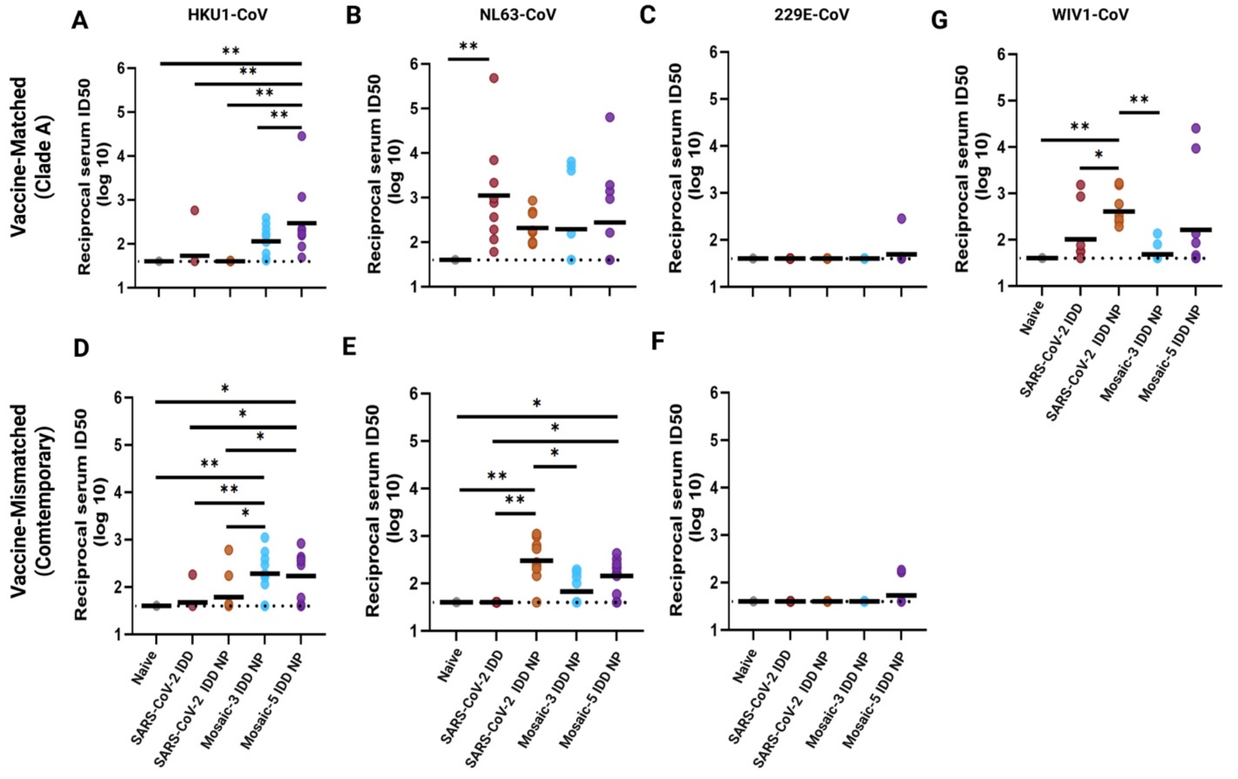


**Extended Data Fig. 5: Comparison of neutralizing antibody response against endemic pseudotyped CoVs and pandemic-threat WIV1-CoV, related to Fig. 4B**. (**A-G**) Sera collected 2-weeks post-boost immunization were evaluated for neutralization of a panel of pseudotyped viruses displaying vaccine-matched (**A-C**) and mismatched (**D-F**): HKU1-CoV, NL63-CoV, 229E spikes and pandemic-threat WIV1-CoV (**G**). The horizontal dashed line represents the limit of detection(LOD). Bar indicates geometric mean of ID50 for N=10 mice per group. Each dot represents a mouse in a group. Two–way ANOVA with Tukey's post-hoc test was performed to convert ID50 values to log10 scale. The statistical significance is displayed as follows: $*p < 0.05$; $**p < 0.01$; $***p < 0.001$; $****p < 0.0001$.

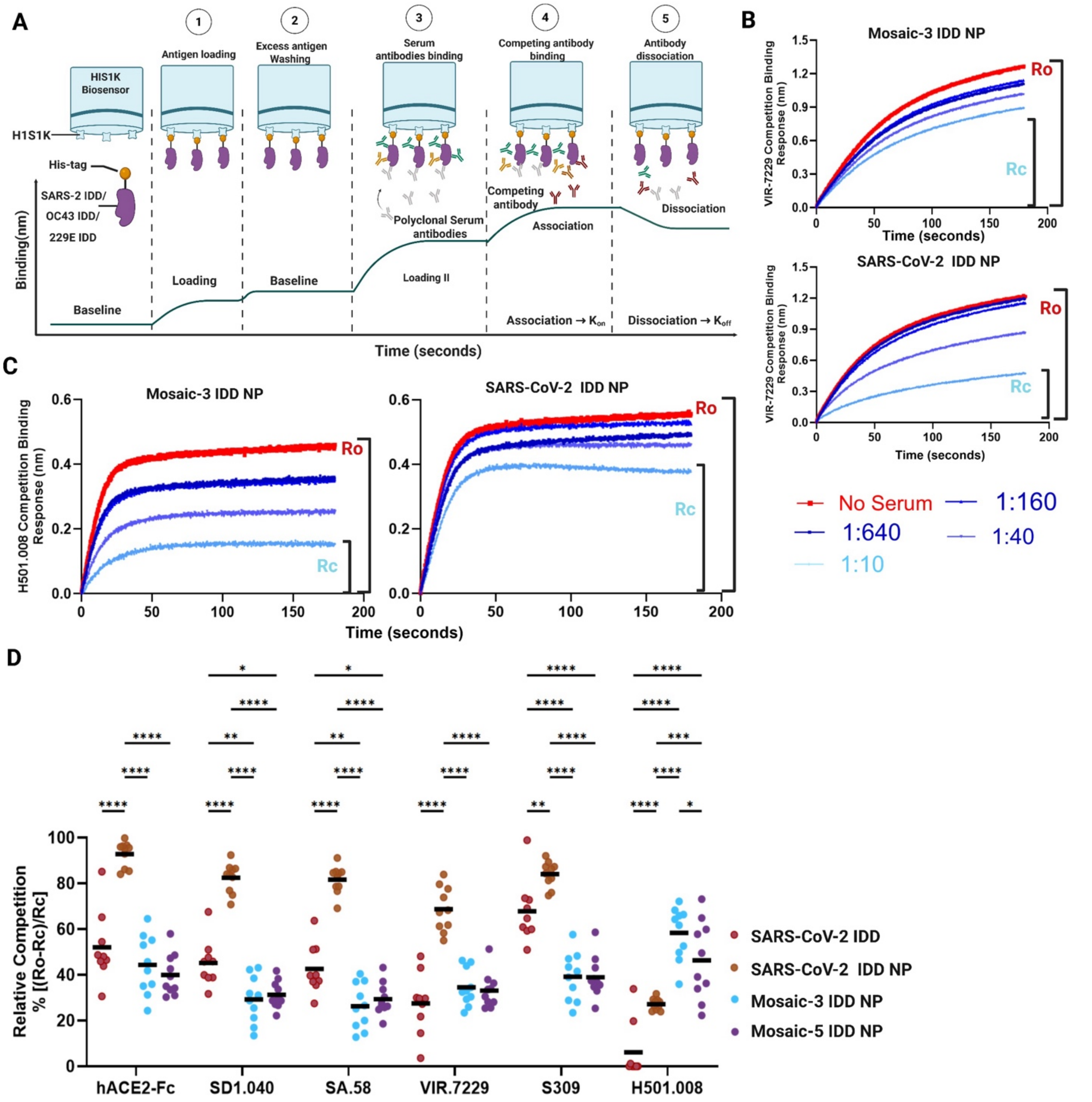

**Extended Data Fig. 6: Detection of antibodies and epitope mapping in immune sera using bio layer interferometry competition binding assay. (A)** Experimental workflow for serum competition assay as measured by BLI. **(B, C)** Representative binding data curves for mAbs VIR-7229 and H501-008 competing with serially diluted immune sera from mice immunized with Mosaic-3 IDD NP and SARS-CoV-2 IDD NP. Ro represents the highest binding signals of VIR-7229 and H501-008 without serum. Rc represents the highest binding signals of VIR-7229 and H501.008 tested with sera from vaccinated mice. (**D**) Cross-competition and inhibition of immune sera by indicated mAbs. 1:10 sera dilution is shown. Bar indicates mean percent inhibition for N=10 mice per group. Each dot represents an individual mouse in a group.

Two–way ANOVA with Tukey's post-hoc test was performed. The statistical significance is displayed as follows: $*p < 0.05$; $**p < 0.01$; $***p < 0.001$; $****p < 0.0001$.

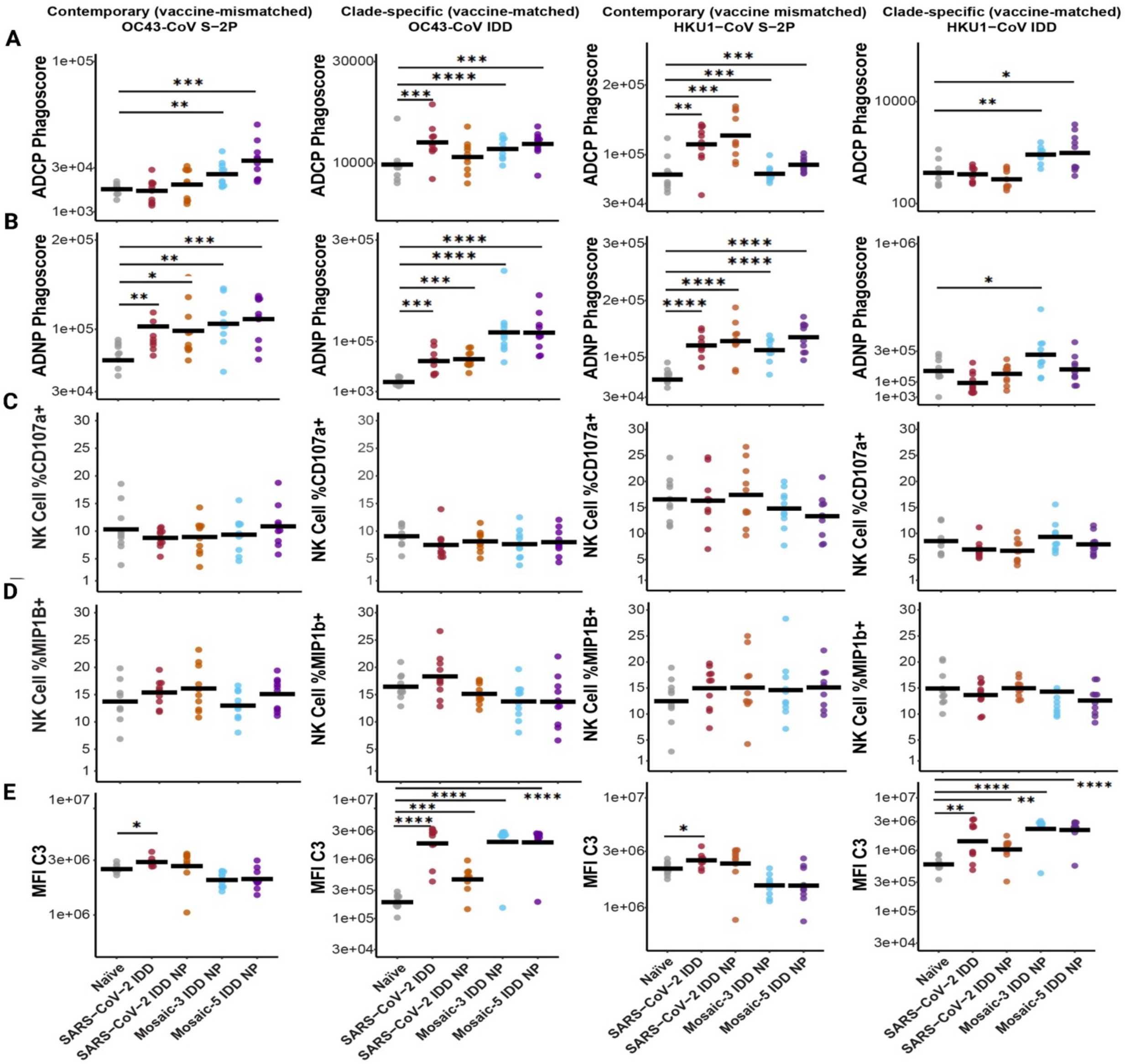


**Extended Data Fig. 7: Fc antibody-dependent functional scores (ADCP, ADNP, ADNK, ADCD) against endemic CoV OC43-CoV and HKU1-CoV.** **(A-E)** Fc-mediated effector functions of the naïve and immune sera after vaccination with IDD NPs against contemporary OC43-COV S-2P (left-hand column), clade-specific OC43-CoV IDD (left-middle column), contemporary HKU1-CoV S-2P (right middle column), and clade-specific HKU1-CoV IDD. Antibody-mediated cellular phagocytosis with monocytes (ADCP, **A**) or neutrophils (ADNP, **B**) using IDD-vaccine-induced immune and naïve sera and beads coated with indicated HKU1-CoV and OC43-CoV proteins. (**C-D)**, Antibody-dependent natural killer cell activation (ADNKA) using IDD-vaccine-induced immune and naïve sera and beads coated with indicated OC43-CoV and HKU1-CoV proteins. **C**, Percentages of NK cells expressing CD107a against the antigens by vaccine group. **D**, Percentages of NK cells expressing MIP-1β against the antigens by vaccine group. The experiment was run with PBMCs from two separate donors. (**E**) Antibody-dependent deposition of complement(ADCD) on beads coated with indicated HKU1-CoV and OC43-CoV proteins and incubated with naïve or immune sera. In all figures, the horizontal black bar indicates the group mean value (N=10 mice per group). Each dot represents an individual mouse in a group. Two–way ANOVA with Tukey's post-hoc test was performed. The statistical significance is displayed as follows: $*p < 0.05$; $**p < 0.01$; $***p < 0.001$; $****p < 0.0001$. Source data file are provided.
