## Supplementary File for "Rational Design of Multiclade Coronavirus Spike Immunodominant Domain Nanoparticles to Elicit Broad Antibody Responses"

### Supplementary Figures

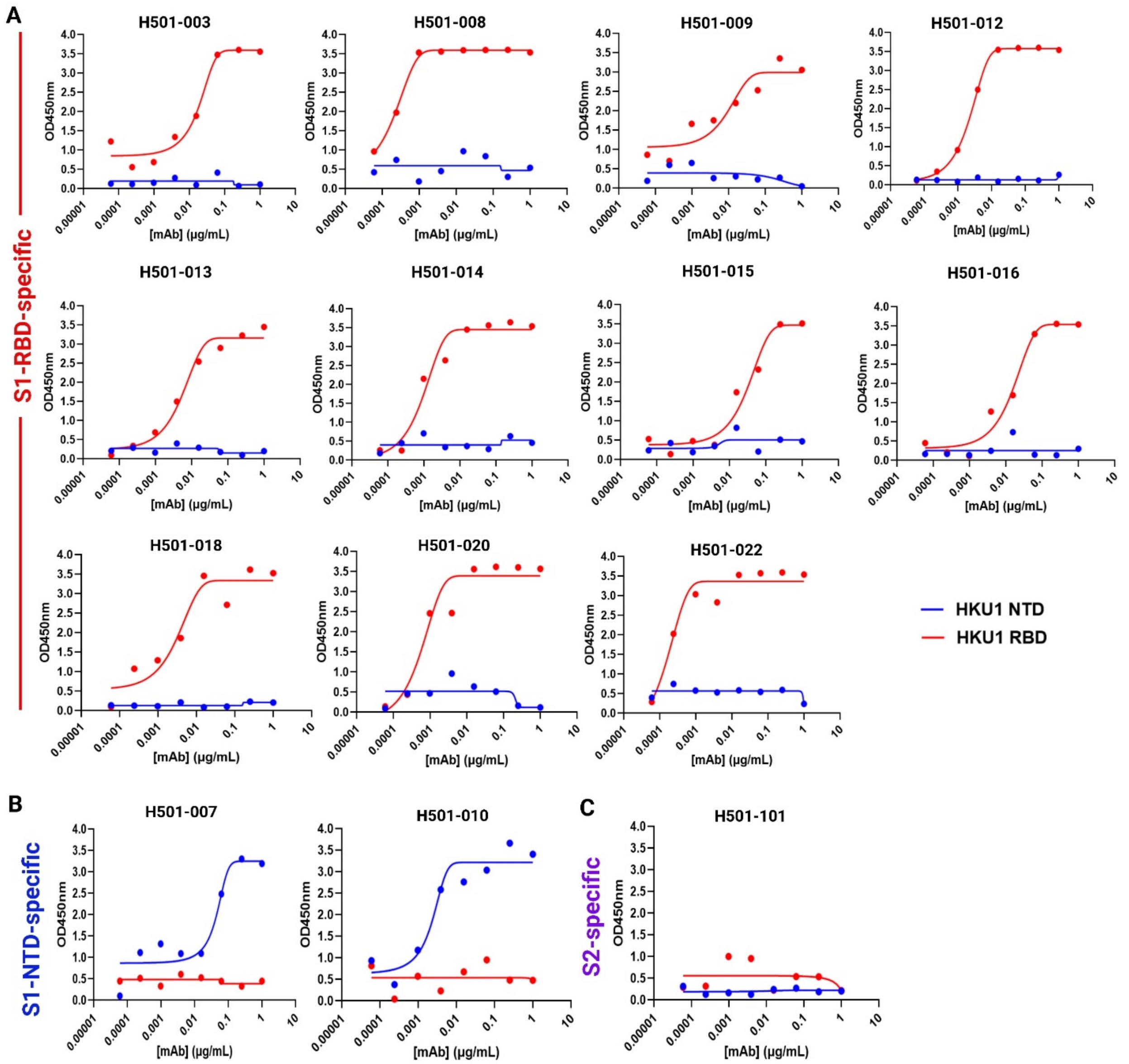


**Supplementary Fig. 1: Antigenic characterization of HCoV-HKU1 S1 domain probes.** (**A**) S1-RBD-specific mAbs, (**B**) S1-NTD-specific mAbs and (**C**) S2-specific mAb isolated from convalescent HCoV-HKU1-positive PBMCs were screened by ELISA for binding to HKU1 RBD (red) and HKU1 NTD (Blue) probes.

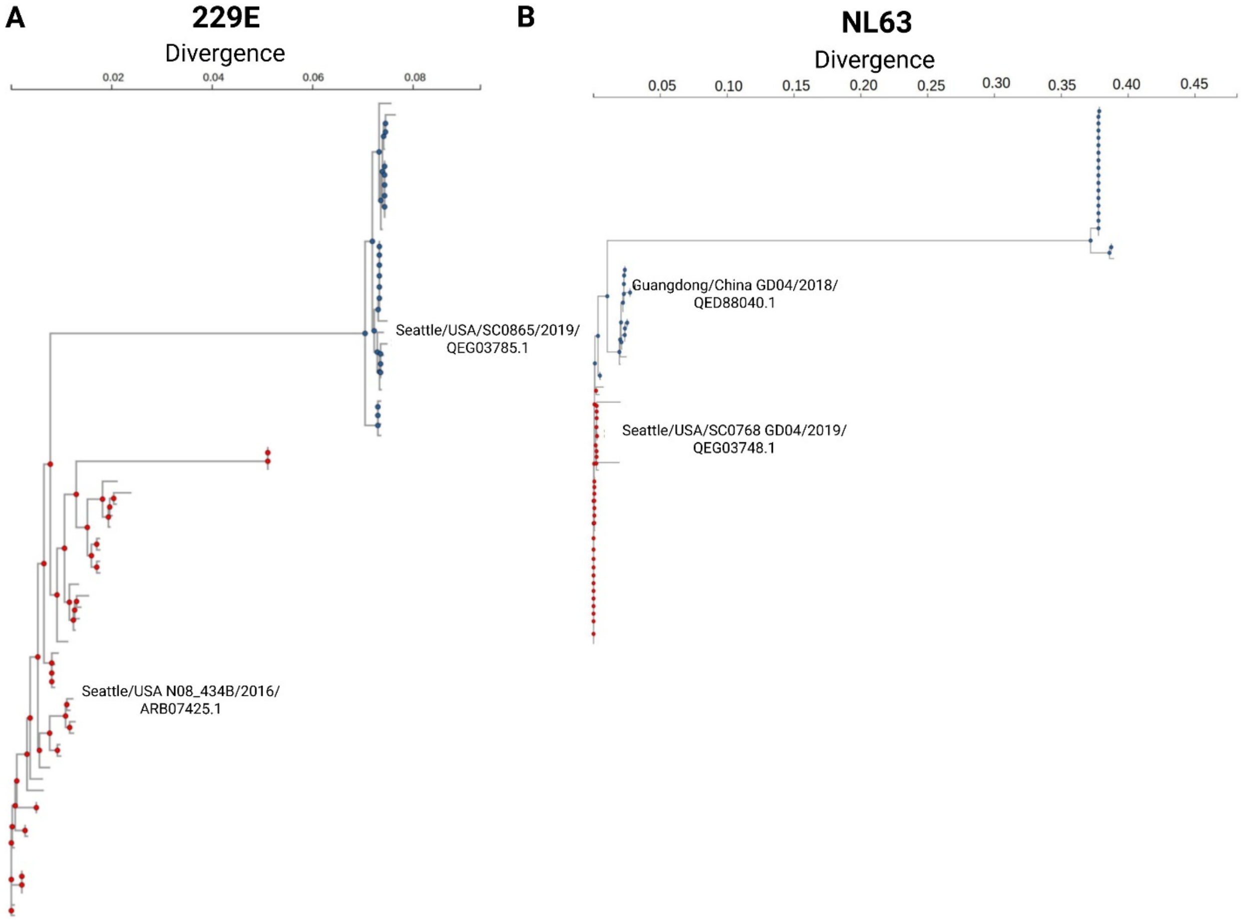
A
229E
Divergence
0.02
0.04
0.06
0.08
Seattle/USA/SC0865/2019/
QEG03785.1
Seattle/USA N08_434B/2016/
ARB07425.1
B
NL63
Divergence
0.05
0.10
0.15
0.20
0.25
0.30
0.35
0.40
0.45
Guangdong/China GD04/2018/
QED88040.1
Seattle/USA/SC0768 GD04/2019/
QEG03748.1

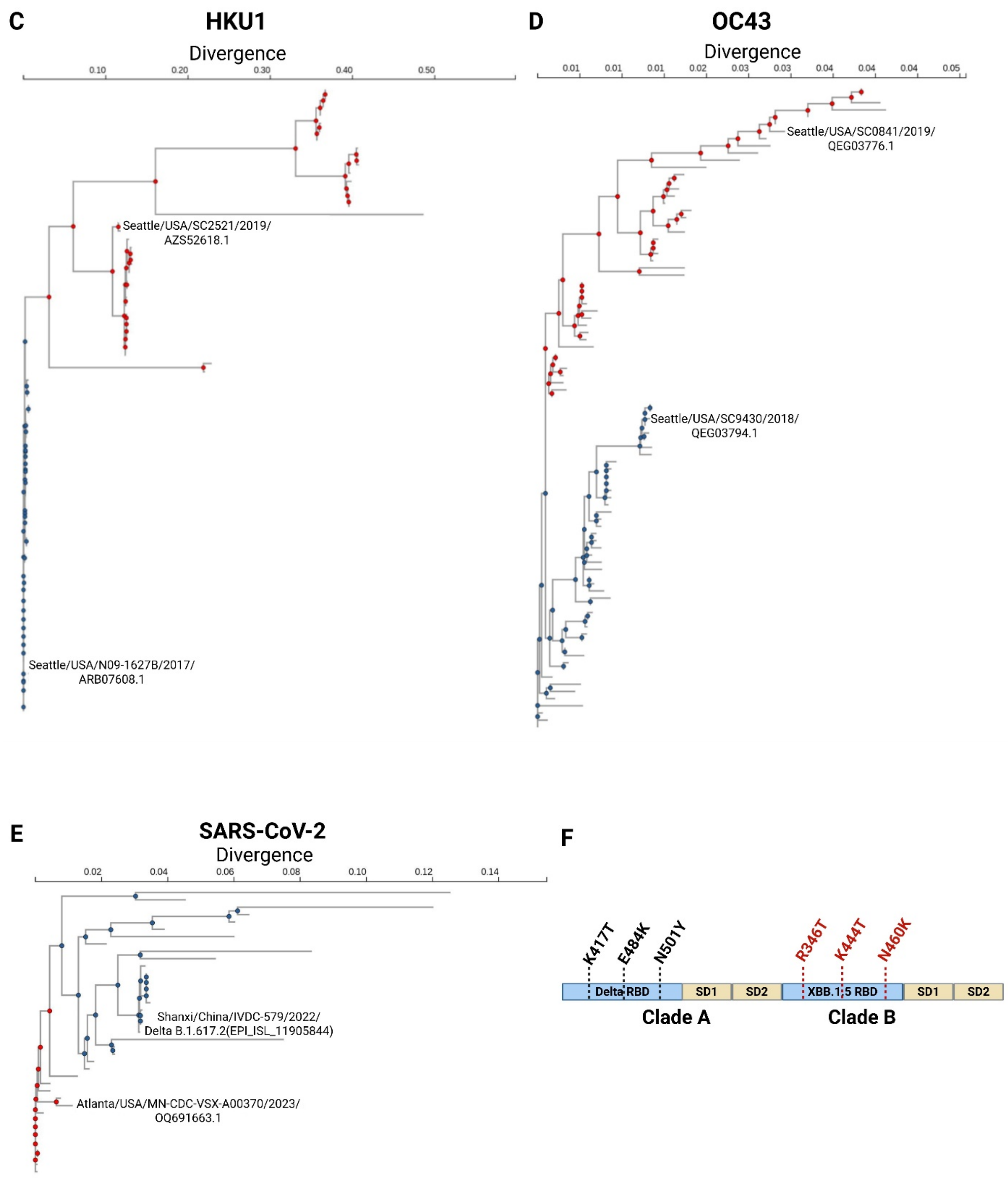


**Supplementary Fig. 2: Selection of EhCoV immunodominant domain (IDD) vaccine antigens, related to Fig. 1H**. Spike Protein Sequences were identified using Nextstrain database and filtered by timeframe. (**A-E**) Phylogenetic trees were created with Clustal Omega EMBL-EBI using seeded guide trees and HMM profile-profile techniques and aligned by Clustal Omega MSA for EhCoV (**A**) 229E, (**B**) NL63 (**C**) HKU1, (**D**) OC43 and (**E**) SARS-CoV-2. Clades A and B are highlighted in blue and red, respectively. Selected S proteins for IDD vaccine designs are indicated. (**F**) Schematic depiction of mutation patches used to expand SARS-CoV-2 cross-variant antigenic coverage on the SARS-CoV-2 IDD construct.

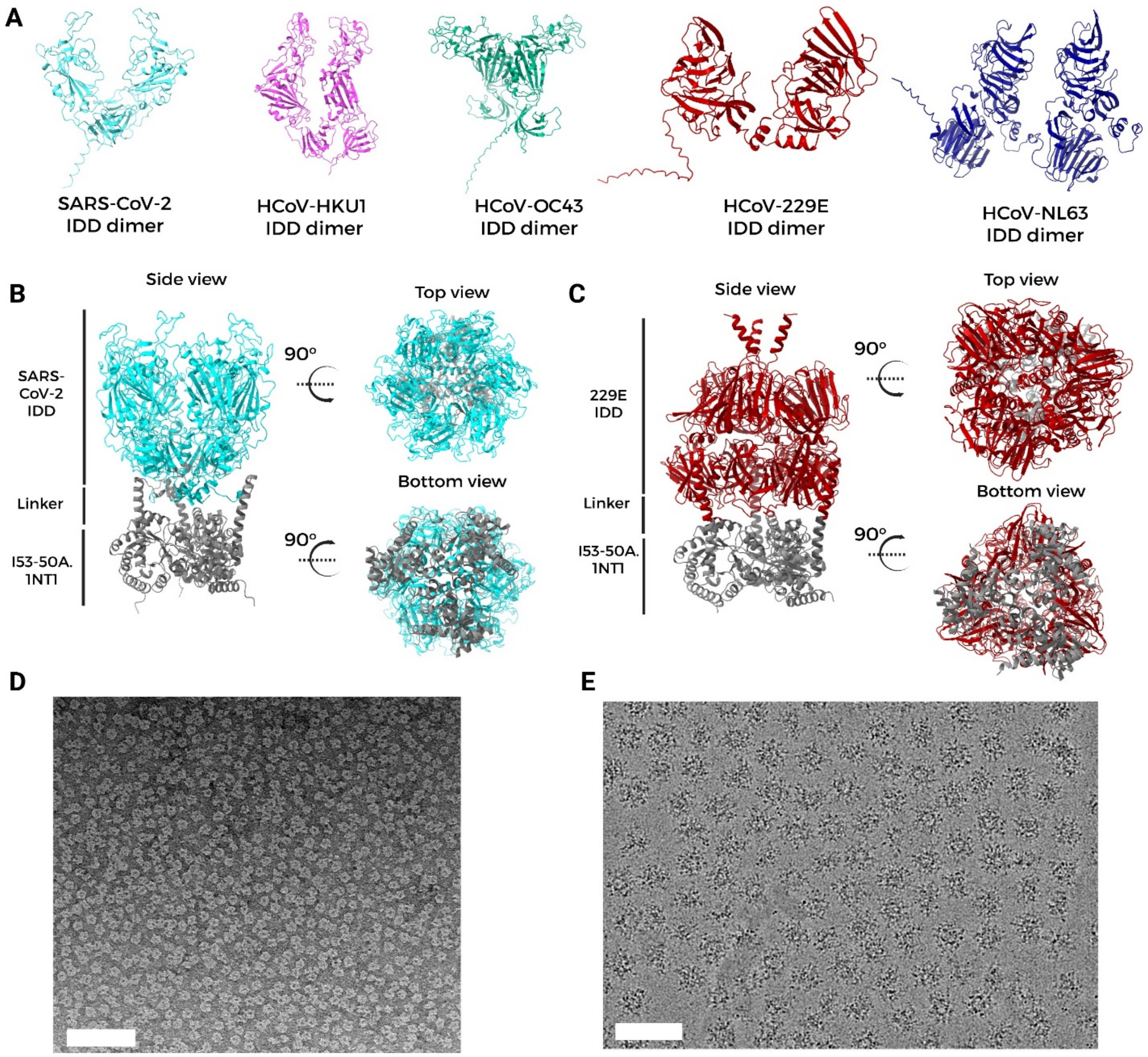


**Supplementary Fig. 3: IDD design and NP characterization, related to Fig. 2.** (**A**) AlphaFold structure prediction of monomeric chimeric IDD antigens. Molecular models of five EhCoV IDDs. (**B-C**) Composite molecular models of chimeric immunodominant domain (IDD) trimers for representative endemic (**B**) β-hCoV (SARS-CoV-2) and (**C**) α-hCoV (229E). The C-terminus of SARS-CoV-2 IDD (cyan) and 229E-CoV IDD (red) were fused to the N-termini of I53-50A trimer (gray) via a linker. (**D**) Representative negative-stain EM of purified bare I53-50 NP. Scale bar, 100 nm. (**E**) Representative cryo-EM micrograph of mosaic-5 IDD NP embedded in vitreous ice. Scale bar, 50 nm.

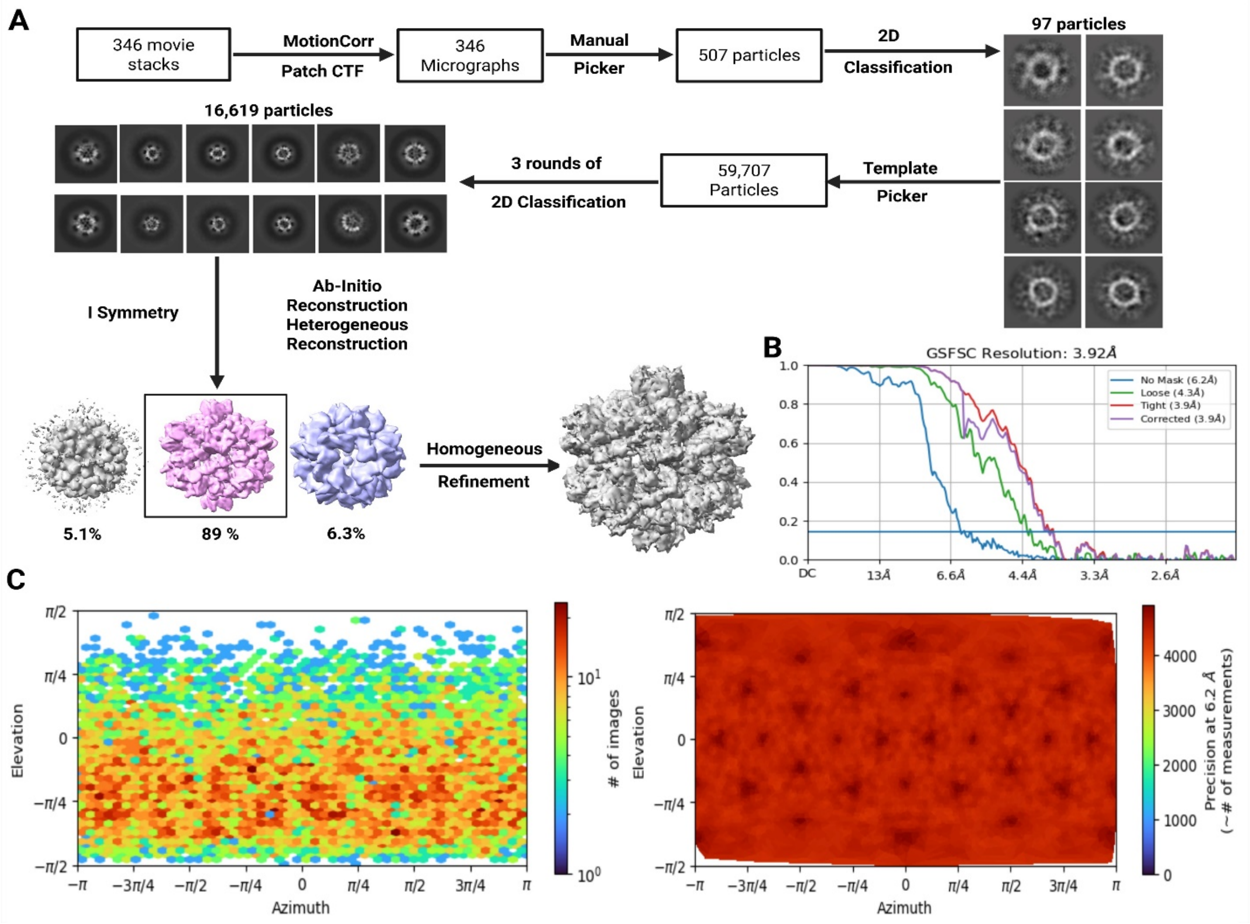


**Supplementary Fig. 4: Cryo-EM structure determination of Mosaic-5 IDD NP, related to Figure 2D-E.** (**A**) Cryo-EM data processing and reconstruction workflow showcasing all relevant and standard steps. 3D reconstruction was done using icosahedral symmetry. Final resolution was 3.9Å. (**B**) Gold-standard Fourier shell correlation (FSC) curve for the Mosaic-5 IDD NP density map shown in **Fig. 2E**. The 0.143 cut-off value is indicated by horizontal blue bar. (**C**) 2D Euler angular distribution plots used in reconstructions.

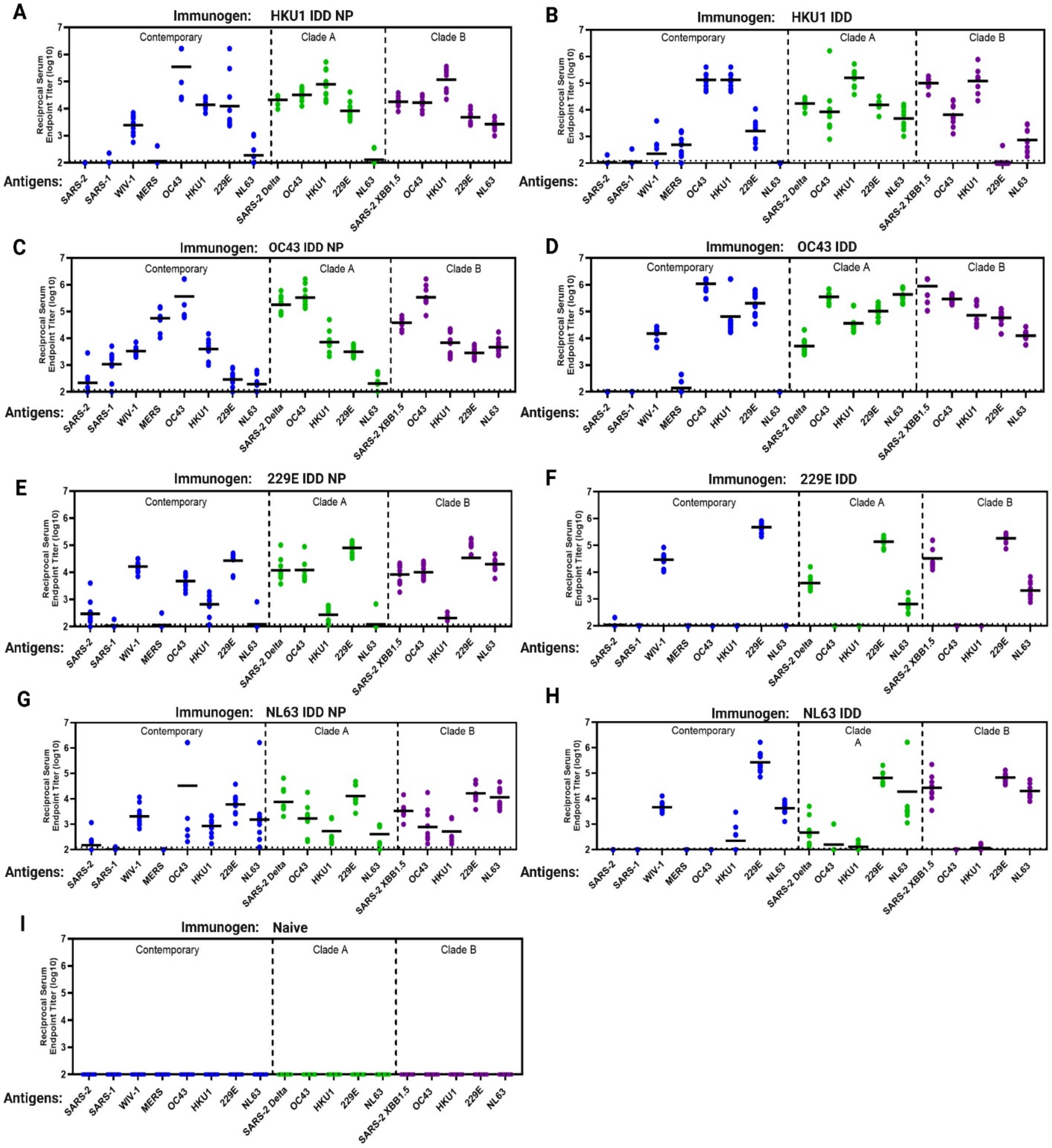


**Supplementary Fig. 5: (A-I) Cross-reactive antibody responses in EhCoV IDD and IDD NP immunized mice, related to Fig. 3**. Female BALB/cJ mice (N = 10/group) were immunized at weeks 0 and 3 with 5 ug of IDD trimer (**A, C, E, G**) or IDD NPs (**B, D, F, H**) or Naive (**I**) with SAS adjuvant and bled at week 5 for serology. Control mice were immunized with adjuvanted PBS. Immune sera were assessed for antibody binding to various CoV S proteins by ELISA. Contemporary CoV S proteins are shown in blue. Clade A CoV S proteins are shown in green. Clade B CoV S proteins are shown in purple. Each dot represents an individual mouse. Horizontal dashed lines represent assay LOD.

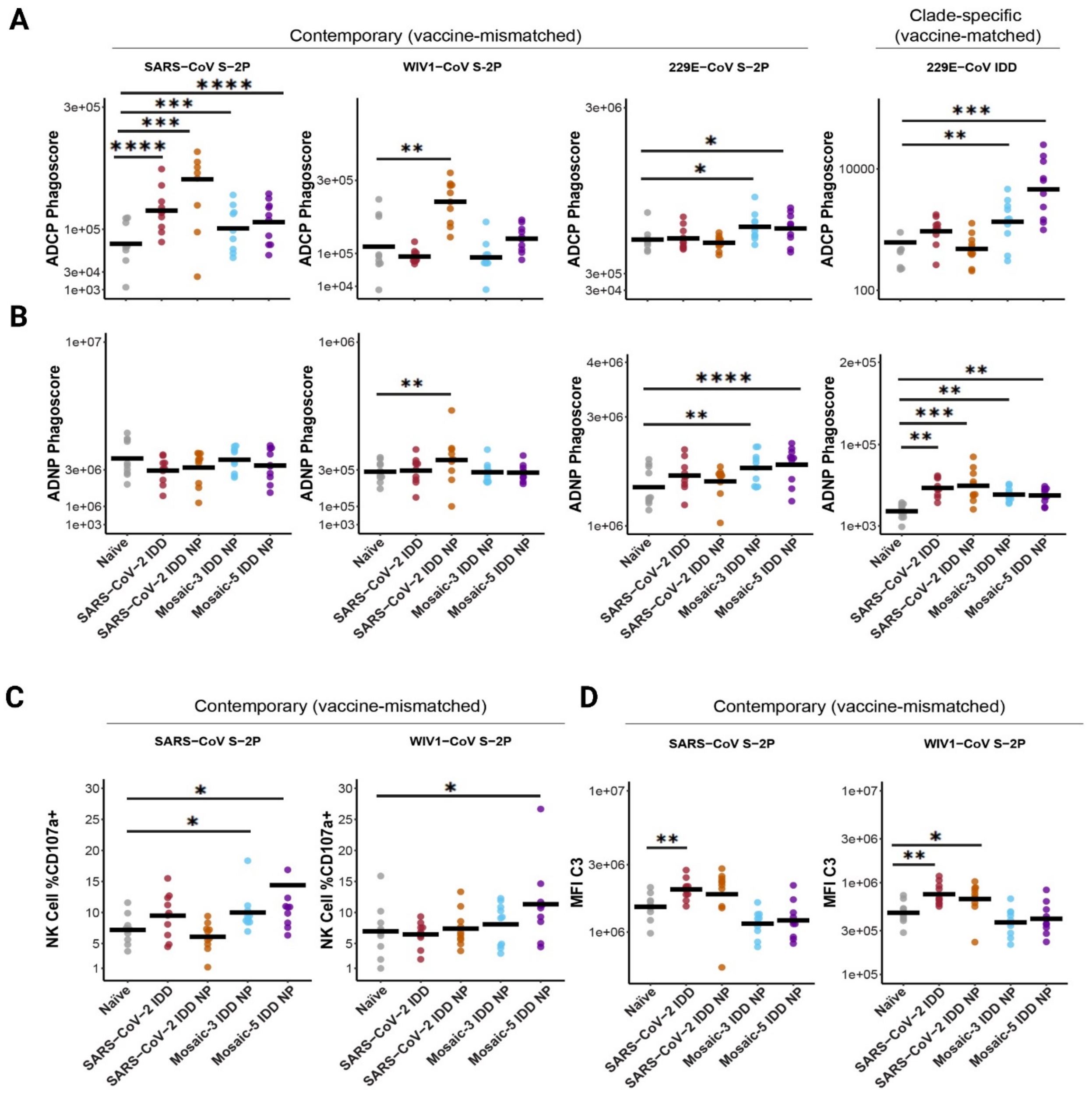


**Supplementary Fig. 6: Antibody Fc-dependent functional scores (ADCP, ADNP, ADNK, ADCD) against pandemic threat and endemic coronavirus(229E-CoV).** (**A, B**) Fc-mediated effector functions of the naïve and immune sera after vaccination with IDD NPs against contemporary (SARS-CoV S-2P, WIV1-CoV S-2P, and 229E-CoV S-2P) and clade-specific (229E-CoV IDD) protein. Antibody-mediated cellular phagocytosis with monocytes (ADCP, **A**) or neutrophils (ADNP, **B**) using IDD-vaccine-induced immune and naïve sera and beads coated with indicated SARS-CoV, WIV1-CoV, and 229E-CoV proteins. (**C**) Antibody-dependent natural killer cell activation (ADNKA) using IDD-vaccine-induced immune and naïve sera and beads coated with indicated SARS-CoV and WIV1-CoV S-2P proteins, as measured by the percentages of NK cells expressing CD107a against the antigens by vaccine group indicated. (**D**) Antibody-dependent complement deposition (ADCD) on beads coated with indicated SARS-CoV and WIV1-CoV S-2P proteins, and incubated with naïve or immune sera. In all figures, the horizontal black bar indicates the group mean value (N=10 mice per group). Each dot represents an individual mouse in a group. Two–way ANOVA with Tukey's post-hoc test was performed. The statistical significance is displayed as follows: $*p < 0.05$; $**p < 0.01$; $***p < 0.001$; $****p < 0.0001$. Source data file is provided.

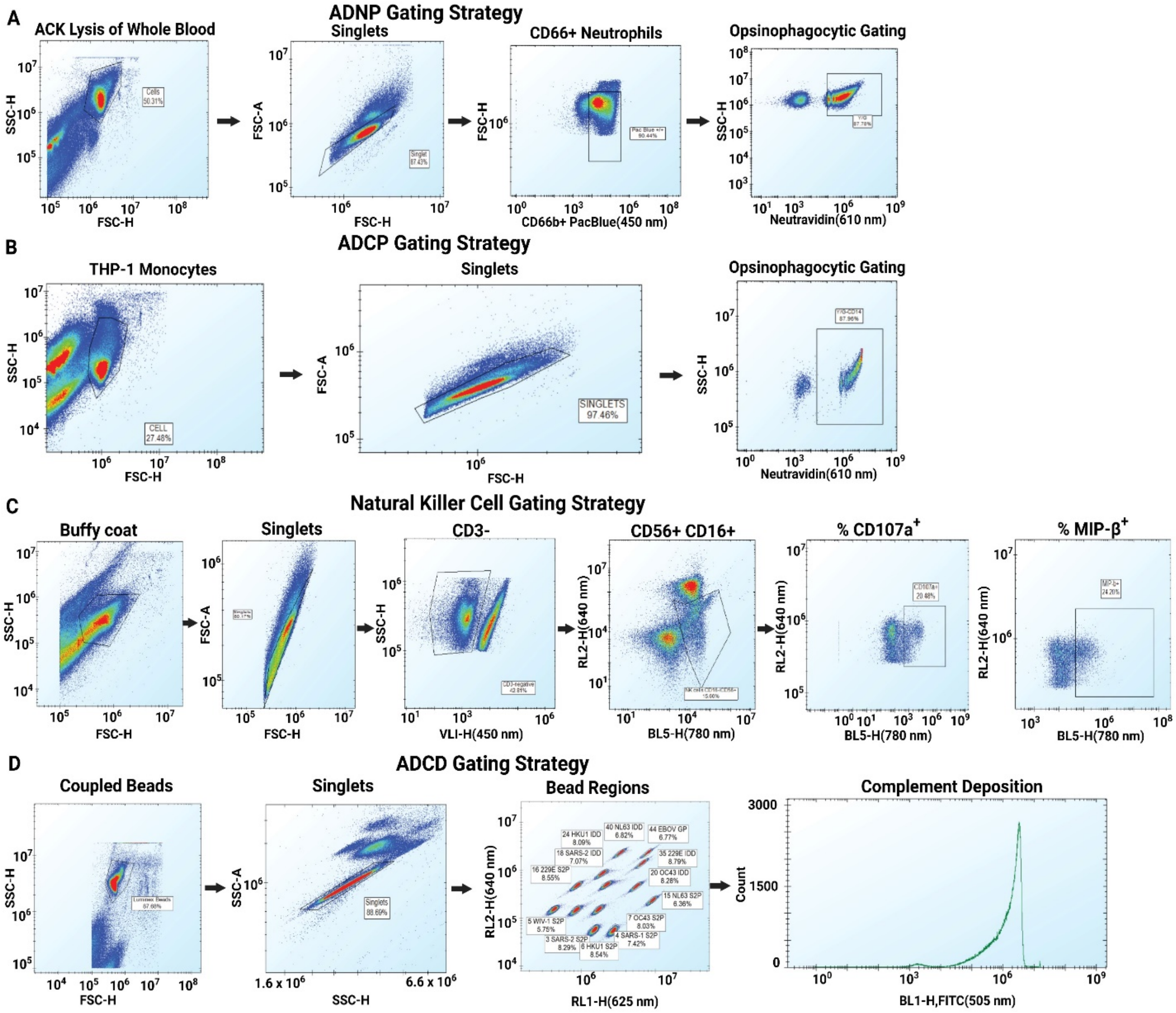


**Supplementary Fig. 7: Flow cytometry gating strategy for Fc effector function assays, related to Fig. 5, Extended data Fig. 7 and Supplementary Fig. 6. (A)** Gating for antibody-dependent neutrophil phagocytosis (ADNP) assay displaying CD66+ neutrophils with phagocytosed beads. (**B**) Gating for antibody-dependent cellular phagocytosis by monocytes (ADCP) assay displaying CD14+ monocytes with phagocytosed beads. (**C**) Gating for antibody-dependent natural killer cell activation (ADNKA) displaying CD56+/CD16-, and CD107a expression. (**D**) Gating for antibody-dependent complement deposition (ADCD) on CoV protein and antibody-coated beads.

### Supplementary Tables

**Supplementary Table. 1**. EhCoV convalescent donor information and sample details

| VRC Study | Subject ID | Date of Confirmed CoV Infection | Sample Type | Blood Draw Information | |
|---|---|---|---|---|---|
| | | | | Date | Months after CoV[1] |
| VRC 200 | A | 1/30/2014 | PBMC | 3/12/2014 | 1 |
| VRC 500 | | | Serum | 2/10/2015 | 13 |
| VRC 317 | B | 5/3/2018 | Serum | 7/11/2018 | 2 |
| | | | PBMC | | |
| | C | 7/12/2018 | Serum | 10/4/2018 | 3 |
| | | | PMBC | | |
| | D | 1/10/2018 | Seum | 4/12/2018 | 3 |
| | | | PBMC | | |
| VRC 200 | E | 3/20/2020 | Serum | 6/24/2020 | 3 |
| | | | PBMC | | |

[1]Date of blood draw – Date of confirmed CoV infection)/12 = Months after CoV

**Supplementary Table .2.** Cryo-EM Data collection and processing parameters, related to Figure 2D-E.

| Cryo-EM Data Collection and Processing Statistics | |
|---|---|
| **EM Data Collection** | |
| Microscope | Talos Arctica |
| Software | SerialEM |
| Voltage(kV) | 200 |
| Detector | Gatan K3 |
| Magnification(Norminal) | 36,000 |
| Electron exposure(e-/ $Å^2$) | 60.04 |
| Defocus range (μm) | 0.1-2.4 |
| Pixel size(Å) | 1.1 |
| Flux (e-/pix/ Å) | 12 |
| Exposure time(seconds) | 4.224 |
| Total frames | 47 |
| Micrographs collected | 346 |

| 3D Reconstruction Statistics | |
|---|---|
| Software | CryoSPARC v4.6.2 |
| Symmetry imposed | I |
| Box size(pix) | 450 |
| Initial particle image no | 346 |
| Final particle image no | 170 |
| Map sharpening B-factor | -134 |
| FSC threshold | 0.143 |
| Unmasked resolution at 0.143FSC(Å) | 6.2 |
| Masked resolution at 0.143FSC(Å) | 3.9 |
| Map resolution(Å) | 3.92 |

**Supplementary Table. 3**: Sequences of designed immunogens and expression constructs reported in this study.

| Construct Name | Amino Acid Sequence |
|---|---|
| **Color Key** | Secretion Signal Sequence, Clade A domain, Clade B domain, GS linker, Display Linker, I53-50A.1NT1, Streptag II, His-8-tag, I53-50B.4PT1 |
| **SARS-CoV-2 IDD-Linker-I53-50A.1NT1** | MRGLGTCLATLAGLLLTAAGRFPNITNLCPFGEVFNATRFASVYAWNRKRISNCVADYSVLYNSASFSTFKCYGVSPTKLNDLCFTNVY<br>ADSFVIRGDEVRQIAPGQTGTIADYNYKLPDDFTGCVIAWNSNNLDSKVGGNYNYRYRLFRKSNLKPFERDISTEIYQAGSKPCNGVK<br>GFNCYF PLQSYGFQPT YGVGYQPYRV VVLSFELLHA PATVCGPKKS TN LVKNKCVN FNFNGLTGTG VLTESNKKFL PFQQFGRDIA<br>DTTDAVRDPQTLEILDITPCSFGGGGSRFPNITNLCPFHEVFNATTFASVYAWNRKRISNCVADYSVIYNFAPFFAFKCYGVSPTKLNDLC<br>FTNVYADSFVIRGNEVSQIAPGQTGNIADYNYKLPDDFTGCVIAWNSNKLDSTPSGNYNYRYRLFRKSKLKPFERDISTEIYQAGNKPC<br>NGVA GPNCYSPLQS YGFRPTYGVG HQPYRVVVLS FELLHAPATV CGPKKSTNLVKNKCVNFNFNGLTGTGVLTE SNKKFLPFQQ<br>FGRDIADTTDAVRDPQTLEILDITPCSFGGGGGGSAEAAAKASSAEAAAKEAAAKEAAAKEEAARKMEELFKKHKIVAVLRANSVEEA<br>IEKAVAVFAGGVHLIEITFTVPDADTVIKALSVLKEKGAIIGAGTVTSVEQCRKAVESGAEFIVSPHLDEEISQFCKEKGVFYMPGVMTPT<br>ELVKAMKLGHDILKLFPGEVVGPEFVKAMKGPFPNVKFVPTGGVDLDNVCEWFDAGVLAVGVGDALVEGDPDEVREKAKEFVEKIR<br>GCTEGSLEWSHPQFEKGSGHHHHHHHH |
| **HCoV-HKU1 IDD-Linker-I53-50A.1NT1** | MDAMKRGLCCVLLLCGAVFVSPSASGSTVKPVATVHRRIPDLPDCDID KWLNNFNVPS PLNWERKIFS NCNFNLSTLL RLVHTDSFSC<br>NNFDESKIYGSCFKSIVLDK FAIPNSRRSD LQLGSSGFLQ SSNYKIDTTS SSCQLYYSLP AINVTINNYN PSSWNRRYGF NNFNLSSHSV<br>VYSRYCFSVNNTFCPCAKPSFASSCKSHKPPSASCPIGTNYRSCESTTVLDHTDWCRCSCLPDPITAYDPRSCSQKKSLVGVGEHCAGFG<br>VDEEKCGVLDGSYNVSCLCSTDAFLGWSYDTCVSNNRCNIFSNFILNGINSGTTCSNDLLQPNTEVYTDVCVDYDLYGITGQGIFKEVS<br>AVYYNSWQNLLYDSNGNIIGFKDFVTNKTYNIFPCYAGGGSTVKPVATVYRRIPNLPDCDIDNWLNNVSVPSPLNWERRIFSNCNFNLS<br>TLLRLVHVDSFSCNNLDKSKIFGSCFNSITVDKFAIPNRRRDDLQLGSSGFLQSSNYKIDTTSSSCQLYYSLPAINVTINNYNPSSWNRRY<br>GFNNFNLSSHSVVYSRYCFSVNNTFCPCAKPSFASSCKSHKPPSASCPIGTKYRHCDLDTTLYVKNWCRCSCLPDPISTYSPNTCPQKKV<br>VVGIGEHCPGLGINEEKCGTQLNHSSCSCSPDAFLGWSFDSCISNNRCNIFSNFIFNGINSGTTCSNDLLYSNTDVSTGVCVNYDLYGITG<br>QGIFKEVSAAYYNDWQNLLYDSNGNIIGFKDFLTNKTYTILPCYSGRVSGGGGSAEAAAKASSAEAAAKEAAAKEAAAKEEAARKME<br>ELFKKHKIVAVLRANSVEEAIEKAVAVFAGGVHLIEITFTVPDADTVIKALSVLKEKGAIIGAGTVTSVEQCRKAVESGAEFIVSPHLDEEI<br>SQFCKEKGVFYMPGVMTPTELVKAMKLGHDILKLFPGEVVGPEFVKAMKGPFPNVKFVPTGGVDLDNVCEWFDAGVLAVGVGDAL<br>VEGDPDEVREKAKEFVEKIRGCTEGSLEWSHPQFEKGSGHHHHHHHH |
| **HCOV-OC43 IDD-Linker-I53-50A.1NT1** | MPMGSLQPLATLYLLGMLVASVLAIADVYRRKPDLPNCNIEAWLNDKSVPSPLNWERKTFSNCNFNMSSLMSFIQADSFTCNNIDAA<br>KIYGMCFSSITIDKFAIPNRRKVDLQLGNLGYLQSSNYRIDTTATSCQLYYNLPAANVSVSRFNPSTWNKRFGFIEDAVFKPQPAGVLT<br>NHDVVYAQHCFKAPKNFCPCSSCSGKNNGIGTCPAGTNSLTCDNLCTLDPITLKAPDTYKCPQSKSLVGIGEHCSGLAVKSDYCGNNS<br>CTCQPQAFLGWSADSCLQGDKCNIFANFILHDVNNGLTCSTDLQKANTEIELGVCVNYDLYGISGQGIFVEVNATYYNSWQNLLYDS<br>NGNLYGFRDYITNRTFMIHSCYSGGGSGVYELNGYTVQPIADVYRRKPDLPNCNIEAWLNDKSVPSPLNWERKTFSNCNFNMSSLMS |

| | |
|---|---|
| | FIQADSFTCNNIDAAKIYGMCFSSITIDKFAIPNGRKVDLQLGNLGYLQSFNYRIDTTATSCQLYYNLPAANVSVSRFNPSTWNKRFGFI<br>EDSVFKPQPAGVLTDHDVVYAQHCFKAPKNFCPCKSNSSLCVGSGPGKNNGIGTCPAGTNYLTCHNLCNPDPITFTGPYKCPQTKSLV<br>GIGEHCSGLAVKSDYCGGNPCTCQPQAFLGWSADSCLQGDKCNIFANLILHDVNSGLTCSTDLQKANTDIKLGVCVNYDLYGISGQGI<br>FVEVNATYYNSWQNLLYDSNGNLYGFRDYITNRTFMIRSCYSGGGGSAEAAAKASSAEAAAKEAAAKEAAAKEEAARKMEELFKK<br>HKIVAVLRANSVEEAIEKAVAVFAGGVHLIEITFTVPDADTVIKALSVLKEKGAIIGAGTVTSVEQCRKAVESGAEFIVSPHLDEEISQFC<br>KEKGVFYMPGVMTPTELVKAMKLGHDILKLFPGEVVGPEFVKAMKGPFPNVKFVPTGGVDLDNVCEWFDAGVLAVGVGDALVEG<br>DPDEVREKAKEFVEKIRGCTEGSLEWSHPQFEKGSGHHHHHHHH |
| **HCoV-229E IDD -Linker-I53-50A.1NT1** | MFVLLVAYALLHIAGCQTTNGTNTSHAVCNGCVGHSENVFAVESGGYIPSNFAFNNWFLLTNTSSVVDGVVRSFQPLLLNCLWSVSGSQ<br>VITGFVYFNGTGRGACKGFYSNASSDVIRYNINFEENLRRGTILFKTSYGAVVFYCTNNTLVSGDAHIPSGTVLGNFYCFVNTTIGNETT<br>SAFVGALPKTVREFVISRTGHFYINGYRYFSLGNVEAVNFNVTNAATTVCTVALASYADVLVNVSQTAIANIIYCNSVINRLRCDQLSGG<br>SQTTNGMNTSHSVCNGCVGHSENVFAVESGGYIPSNFAFNNWFLLTNTSSVVDGVVRSFQPLLLNCLWSVSGSRLTTGFVYFNGTGRG<br>DCKGFYSNASSDVIRYNINFEENLRRGTILFKTSYGAVVFYCTNNTLVSGDAHIPSGTVLGNFYCFVNTTIGNETTSAFVGALPKTVREF<br>VISRTGHFYINGYRYFSLGNVEAVNFNVTNAATTVCTVALASYADVLVNVSQTAIANIIYCNSVINRLRCDQLSGGGGSAEAAAKASSA<br>EAAAKEAAAKEAAAKEEAARKMEELFKKHKIVAVLRANSVEEAIEKAVAVFAGGVHLIEITFTVPDADTVIKALSVLKEKGAIIGAGTV<br>TSVEQCRKAVESGAEFIVSPHLDEEISQFCKEKGVFYMPGVMTPTELVKAMKLGHDILKLFPGEVVGPEFVKAMKGPFPNVKFVPTGG<br>VDLDNVCEWFDAGVLAVGVGDALVEG DPDEVREKAKEFVEKIRGCTEGSLEWSHPQFEKGSGHHHHHHHH |
| **HCoV-NL63 IDD-Linker-I53-50A.1NT1** | MKLFLILLILPLVSCFSTCNSNASISMLQLGVPDNSSTIVTGLLPVHWICANQSTSTYPANGFFYIDVGKHRSAFALHSG YYDANQYYIY<br>LTNNISLNAPVTLKICKFGNTSFACLSNVSTSHHCIVNSSFTEQLGVPLGITISGETVRLHLYNATRTFYVPAAYKLTKLSVKCYFSESC<br>VFSVVNATVTVNVTTLNGRIVNYTVCDDCNGYTDNIFSVQQDGRIPNGFSFNNWFLLTNGSTLVDGVSRLYQPLRLTCLWPVPGLKSS<br>TGFVYFNATGSDVNCNGYQHHSVADVMRYNLNFSSNSVDNLKSGVIVFKTLQYDVLFYCSNSSSGVLDTTIPFGPSSQPYYCFINSTIN<br>TTHVSTFVGVLPPIVREIVVARTGQFYINGFKYFDLGFIEAVNFNVTTASATDFWTVAFATFVDVLVNVSATNIQNLLYCDSPFEKLQC<br>EHLQFGLQDGFYSANFLDDNVLPDGGSSNANLSMLQLGVPDNSSTIVTGLLPTHWICANQSTSVYSANGFFFIDVGNHRSAFALHTG<br>YYDVNQYYIYVTNEIGLNASVTLKICKFGNTTFDFLSNSSSSFDCIVNLLFTEQLGAPLGITISGETVRLHLYNVTRTFYVPAAYKLTKL<br>SVKCYFNYSCVFSVVNATVTVNVTTHNGRVVNYTVCDDCNGYTDNIFSVQQDGRIPNGFPFNNWFLLTNGSTLVDGVSRLYQPLRLT<br>CLWPVPGLKSSTGFVYFNATGSDVNCNGYQHNSVADVMRYNLNFSANSVDNLKSGVIVFKTLQYDVLFYCSNSSSGVLDTTIPFGPSS<br>QPYYCFINSTINTTHVSTFVGVLPPTVREIVVARTGQFYINGFKYFDLGFIEAVNFNVTTASATDFWTVAFATFVDVLVNVSATKIQNLL<br>YCDSPFEKLQCEHLQFGLQDGFYSANFLDDNGGGGSAEAAAKASSAEAAAKEAAAKEAAAKEEAARKMEELFKKHKIVAVLRANS<br>VEEAIEKAVAVFAGGVHLIEITFTVPDADTVIKALSVLKEKGAIIGAGTVTSVEQCRKAVESGAEFIVSPHLDEEISQFCKEKGVFYMPG<br>VMTPTELVKAMKLGHDILKLFPGEVVGPEFVKAMKGPFPNVKFVPTGGVDLDNVCEWFDAGVLAVGVGDALVEGDPDEVREKAKE<br>FVEKIRGCTEGSLEWSHPQFEKGSGHHHHHHHH |
| **I53-50A.1NT1** | MKMEELFKKHKIVAVLRANSVEEAIEKAVAVFAGGVHLIEITFTVPDADTVIKALSVLKEKGAIIGAGTVTSVEQCRKAVESGAEFIVS<br>PHLDEEISQFCKEKGVFYMPGVMTPTELVKAMKLGHDILKLFPGEVVGPEFVKAMKGPFPNVKFVPTGGVDLDNVCEWFDAGVLAV<br>GVGDALVEG DPDEVREKAKEFVEKIRGCTEGSLEHHHHHHHH |
| **I53-50B.4PT1** | MNQHSHKDHETVRIAVVRARWHAEIVDACVSAFEAAMRDIGGDRFAVDVFDVPGAYEIPLHARTLAETGRYGAVLGTAFVVNGGIY<br>RHEFVASAVINGMMNVQLNTGVPVLSAVLTPHNYDKSKAHTLLFLALFAVKGMEAARACVEILAAREKIAAGSLEHHHHHHHH |
